# Different low-complexity regions of SFPQ play distinct roles in the formation of biomolecular condensates

**DOI:** 10.1101/2022.11.30.518278

**Authors:** Andrew C. Marshall, Jerry Cummins, Simon Kobelke, Tianyi Zhu, Jocelyn Widagdo, Victor Anggono, Anthony Hyman, Archa H. Fox, Charles S. Bond, Mihwa Lee

## Abstract

Demixing of proteins and nucleic acids into condensed liquid phases is rapidly emerging as a ubiquitous mechanism governing the organisation of molecules within the cell. Long disordered low complexity regions (LCRs) are a common feature of proteins that form biomolecular condensates. RNA-binding proteins with prion-like composition have been highlighted as drivers of liquid demixing to form condensates such as nucleoli, paraspeckles and stress granules. Splicing factor proline- and glutamine-rich (SFPQ) is an RNA- and DNA-binding protein essential for DNA repair and paraspeckle formation. Here, we show that the shorter C-terminal LCR of SFPQ is the main region responsible for the condensation of SFPQ *in vitro* and in the cell. In contrast, we find that, unexpectedly, the longer N-terminal prion-like LCR of SFPQ attenuates condensation, suggesting a more regulatory role in preventing aberrant condensate formation in the cell. Our data add nuance to the emerging understanding of biomolecular condensate formation, by providing the first example of a common multifunctional nucleic acid-binding protein with an extensive prion-like region that serves to regulate rather than drive condensate formation.

**Graphical Abstract:** 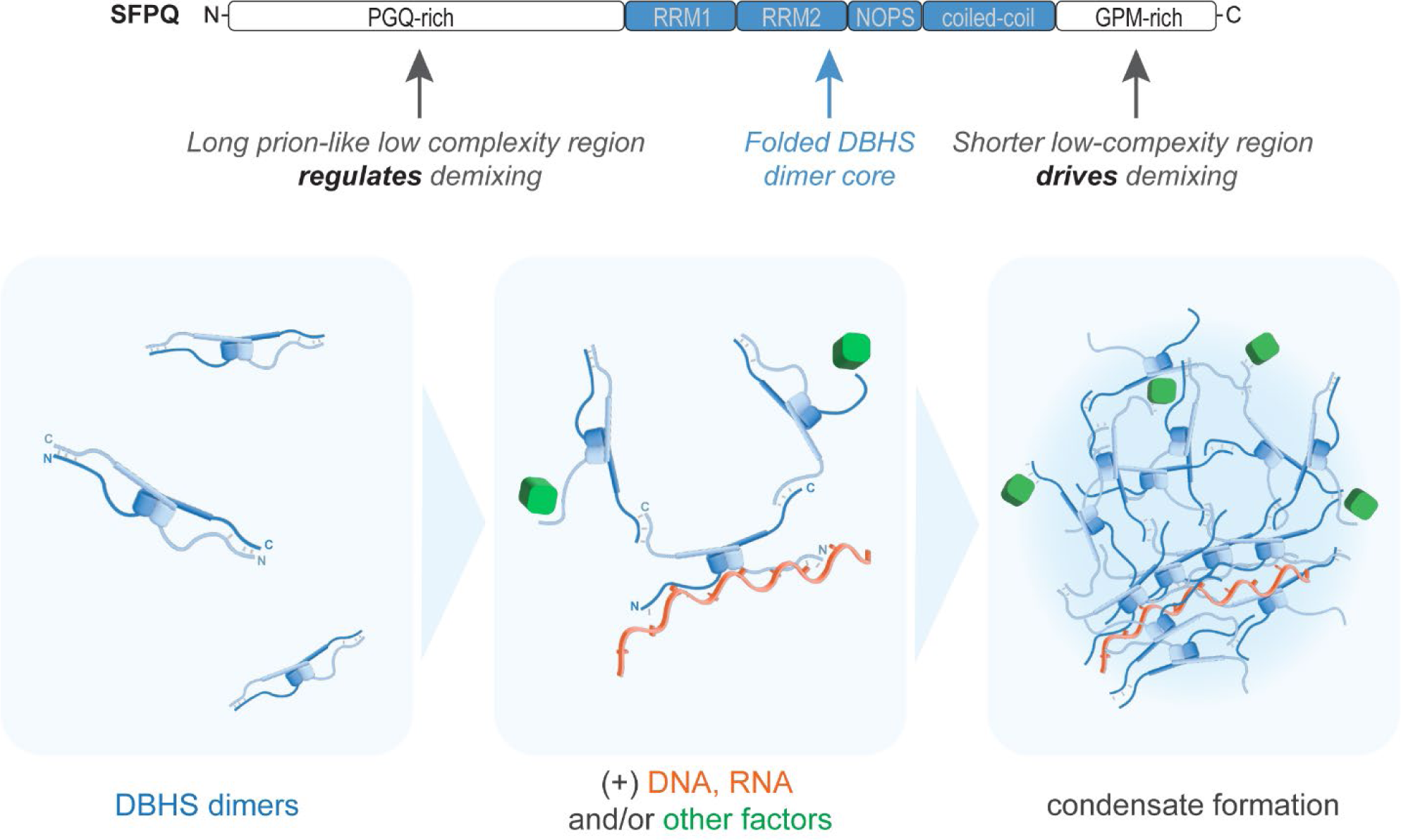

## INTRODUCTION

Protein low-complexity regions (LCRs) are most simply defined as sequences enriched in only a few amino acids and are commonly intrinsically disordered. Enormous interest in disordered LCRs has arisen in recent years due to the observation that they are commonly associated with the condensation of proteins and nucleic acids to form distinct membraneless cellular bodies with defined molecular compositions and physical properties. Biomolecular condensates such as the nucleolus, Cajal bodies, transcriptional hubs, nuclear speckles and paraspeckles are central to the spatial and temporal regulation of biochemical reactions in the mammalian cell nucleus (Banani *et al*, 2017; McCluggage & Fox, 2021; Wei *et al*, 2020). Mounting evidence suggests that the formation of these condensed bodies can be readily described using the soft matter physics concepts of phase separation (Feric *et al*, 2016), microphase separation (Yamazaki *et al*, 2021) and surface wetting (Morin *et al*, 2022). The amino acid sequence and composition of LCRs have been shown to be critical for determining the condensation propensity of a protein and maintaining the reversible liquid-like state of the condensate (Borcherds *et al*, 2021; Bremer *et al*, 2022; Martin & Mittag, 2018; Molliex *et al*, 2015; Wang *et al*, 2018). Although the establishment of rules relating sequence to material state is still in its infancy, various sequence features of LCRs have been shown to facilitate intermolecular interactions important for driving liquid demixing/condensation. These features are often overlapping and include prion-like domains (PLDs) (Bremer *et al*., 2022; Guenther *et al*, 2018), low-complexity aromatic-rich kinked segments (LARKS) (Hughes *et al*, 2018), charge-patterning (Nott *et al*, 2015) and regions rich in basic and aromatic residues (Wang *et al*., 2018). These sequence features are over-represented in nucleic acid-binding proteins involved in subnuclear organisation and regulation of gene expression (Hennig *et al*, 2015; March *et al*, 2016). In addition, both RNA and DNA have been suggested to provide multivalent interactions to drive condensation (Fox *et al*, 2018; Morin *et al*., 2022; Wei *et al*., 2020; Zhang *et al*, 2015), providing a conceptual framework for understanding the formation of condensates containing proteins and nucleic acids.

Splicing factor proline and glutamine-rich (SFPQ; previously known as PSF) is an abundant nuclear DNA- and RNA-binding protein with essential roles in transcriptional elongation, mRNA processing and modification, and DNA repair (Lim *et al*, 2020; Yarosh *et al*, 2015). SFPQ is also an essential component of subnuclear bodies called paraspeckles, which are biomolecular condensates shown to be important for cellular stress response in the contexts of viral infection, cancer, early development, and neurodegeneration (Fox *et al*., 2018; McCluggage & Fox, 2021). Paraspeckle abundance scales with the level of NEAT1_2 long noncoding RNA (Hirose *et al*, 2014). SFPQ plays an essential role in determining NEAT1_2 levels by binding and protecting the RNA (Sasaki *et al*, 2009; West *et al*, 2016). Paraspeckles can either be singular spheroids of 360 nm diameter, or chains of spheroids with a constant width of 360 nm, and length up to 1.5 μm (Souquere *et al*, 2010; West *et al*., 2016). In humans, SFPQ has two paralogues: NONO and PSPC1. They are all members of the ‘*Drosophila*-Behaviour, Human Splicing’ (DBHS) protein family, and exist as obligate homo- or heterodimers in the nucleus (Knott *et al*, 2016). DBHS proteins are composed of a central globular region flanked on either side by long LCRs (**Figure 1a**). The central region, which is highly conserved between DBHS family members, contains tandem RNA-recognition motif (RRM) domains followed by a unique NonA/paraspeckle (NOPS) domain and an extended coiled-coil region (Knott *et al*, 2015). Multiple in-depth structural studies have shown that these domains pack together to form a compact, well-ordered DBHS core, which is essential for RNA-binding and protein dimerisation (**Figure 1c**) (Huang *et al*, 2018; Knott *et al*, 2022; Lee *et al*, 2022; Passon *et al*, 2012). In addition, the extended coiled-coil drives the polymerisation of SFPQ dimers, which is critical for the functions of SFPQ in transcriptional regulation and subnuclear body formation (Lee *et al*, 2015). In contrast, the flanking LCRs of DBHS proteins are highly divergent and remain largely unstudied. Phylogenetic analysis suggests that, of the three DBHS paralogues, SFPQ is the most divergent from their common ancestral DBHS protein, and that this is largely due to low selection pressure within its conspicuously longer N-terminal LCR (Knott *et al*., 2015).

**Figure 1.**
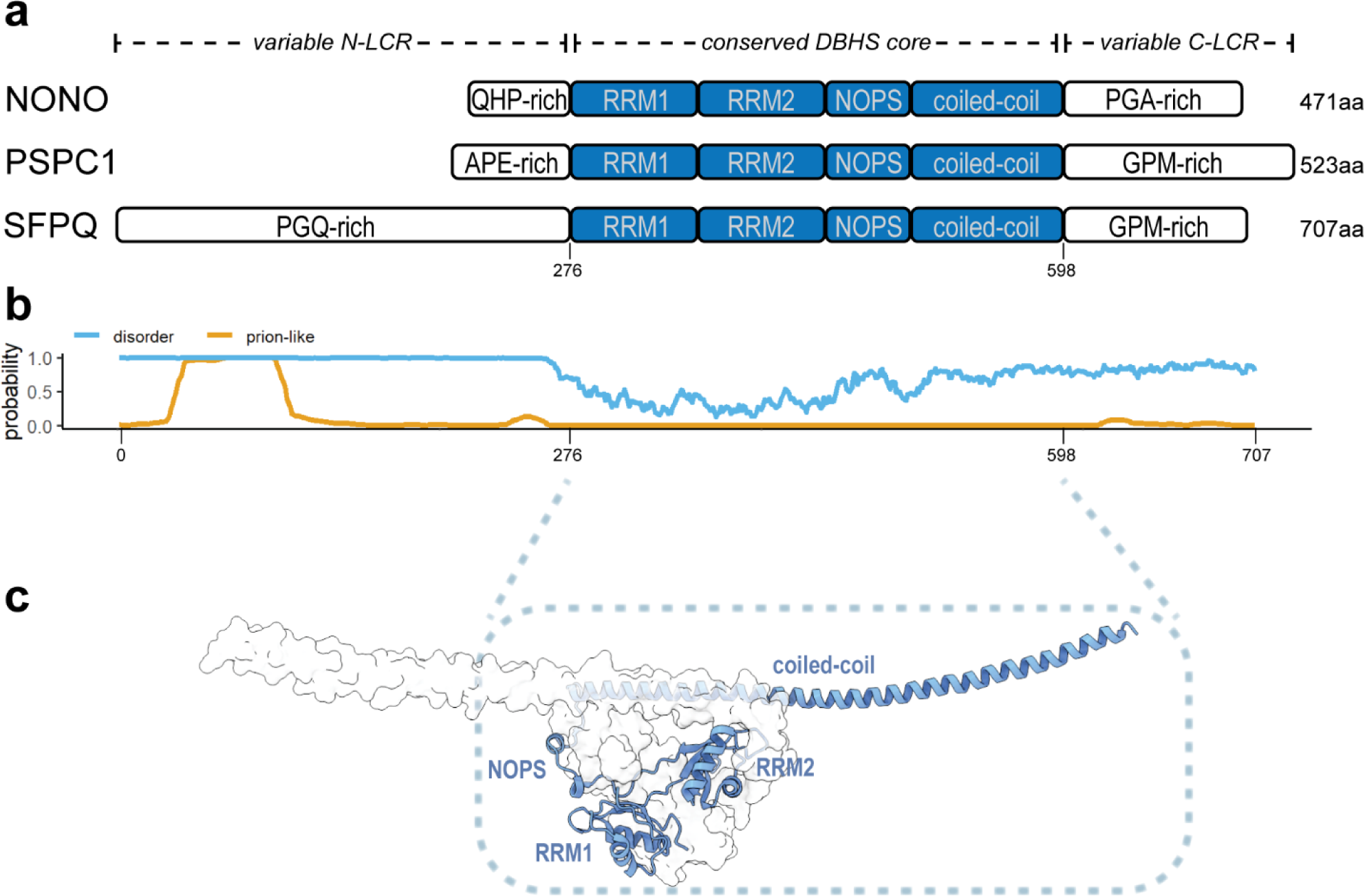
SFPQ is a DBHS protein with an extended N-terminal prion-like low-complexity region. **(a)** Schematic showing domain topologies of the three human DBHS proteins. The well-ordered folded domains of the DBHS core (blue) are flanked on either side by variable regions (white) enriched in specific amino acids, as indicated. Relative lengths of each region are shown to scale, highlighting the extended N-LCR of SFPQ. The total number of amino acids in each protein is indicated on the right. **(b)** *In silico* prediction of intrinsically disordered regions (IUPred2A; blue) and regions that are prion-like (PLAAC; orange) within SFPQ suggest that both LCRs are disordered, and that a large section of the N-LCR is prion-like. **(c)** Crystal structure of the homodimer formed by the core conserved DBHS region of SFPQ (PDB 4wij). Structural domains of one dimeric partner (shown in cartoon representation) are labelled. The other dimeric partner is shown in surface representation.

Using *IUPred2A* (Erdős & Dosztányi, 2020) we find that both LCRs of SFPQ, which flank the globular conserved DBHS region, are predicted to be intrinsically disordered (**Figure 1b**). However, each is very different in length and displays a different amino acid enrichment profile. The shorter C-terminal LCR (C-LCR) of SFPQ is similar in length to the corresponding regions of NONO and PSPC1 (∼100 amino acids), and is enriched in glycine, proline and methionine residues. The N-terminal LCR (N-LCR) is approximately 275 amino acids long, is enriched in glycine, proline and glutamine residues, and is characterised by multiple poly-Q and poly-P tracts. The length of the N-LCR of SFPQ is the most obvious feature distinguishing SFPQ from NONO and PSPC1, which contain N-LCRs of only 50-60 amino acids (**Figure 1a**). Using the *PLAAC* algorithm (Lancaster *et al*, 2014) we find that a large proportion of the SFPQ N-LCR is predicted to have a “prion-like” amino acid composition (**Figure 1b**), suggesting a role in facilitating protein-protein interactions that may be important for driving condensation of the protein (Bremer *et al*., 2022; Hennig *et al*., 2015; Peran & Mittag, 2020).

Here, we have used *in vitro* assays, fluorescence cell imaging and DNA damage recruitment assays to investigate the roles of the N- and C-LCRs of SFPQ in driving protein condensation. Unexpectedly, we find that the shorter C-LCR of SFPQ is the main driver of condensation, whereas the extended prion-like N-LCR attenuates condensation of the full-length protein. Taken together these data show that the different LCRs of SFPQ have different and possibly opposing roles in determining the material state, subnuclear localisation and functions of SFPQ inside the cell.

## MATERIALS AND METHODS

### Plasmids for heterologous protein expression

For producing proteins of interest in *Escherichia coli* for *in vitro* experiments, an open reading frame (ORF) for full-length SFPQ protein (residues 1-707; reference sequences: NCBI NM_005066.3 / Genbank X70944.1 / UniProt P23246-1) with an N-terminal 6×His-mEGFP-TEV tag was inserted using Gibson assembly between the ribosome binding site and T7 terminator of pET-28a(+) (Novagen). Expression vectors for H6-mEGFP-SFPQ ΔN-LCR (SFPQ residues 276-707), ΔC-LCR (residues 1-598) and ΔN&ΔC (residues 276-598) truncations were then constructed from this plasmid by site-directed deletion of the appropriate regions of the SFPQ ORF using the Q5® Site-Directed Mutagenesis Kit (New England Biolabs). The mammalian expression construct encoding GFP-tagged full-length SFPQ was described previously (Huang *et al*, 2020; Lee *et al*., 2015). The truncated constructs [pEGFP-SFPQ ΔN-LCR (residues 276-707), pEGFP-SFPQ ΔC-LCR (residues 1-598 plus 700-707), and pEGFP-SFPQ ΔN- and ΔC-LCR (residues 276-598 plus 700-707)] were generated by site-directed deletion using the Q5 site-directed mutagenesis method (New England Biolabs). To ensure nuclear localisation of the truncated constructs, the endogenous nuclear localisation signal (NLS) of SFPQ (residues 700-707) was kept in ΔC-LCR constructs. All constructs were verified by DNA sequencing.

### Protein expression and purification for *in vitro* experiments

Plasmids for IPTG-inducible over-expression of H6-mEGFP-SFPQ proteins were transformed into *E. coli* Rosetta™ 2(DE3) cells (Novagen). Transformed cells were cultured overnight at 37°C in lysogeny broth (LB) with selection (50 μg/mL kanamycin, 34 μg/mL chloramphenicol). 30 mL of the overnight culture was then added to 3 L of fresh LB + 50 μg/mL kanamycin + 17 μg/mL chloramphenicol in six 2.5 L conical flasks and incubated at 37°C, shaking (180 rpm). Protein expression was induced at OD_600_ ∼ 0.5 by adding 0.5 mM IPTG, before continuing growth for 18-20 h at 20°C. All subsequent purification steps were performed at ambient temperature. Cells were harvested and resuspended in 100 mL of lysis buffer (1 M KCl, 5% glycerol, 10 mM imidazole, 50 mM Tris-HCl, 250 mM L-arginine, 1 mM PMSF, 0.05 mg DNase I) per L of cell culture. Cells were then lysed using either an EmulsiFlex-C5 high pressure homogenizer (Avestin) at ∼ 15,000 psi, or by sonication on ice (cycles of 7 sec on, 3 sec off, for a total on-time of 6 min; 30% amplitude). Clarified lysate was loaded onto a 5 mL Ni^2+^-charged HisTrap HP column (Cytiva), washed with 10 column volumes (CV) of binding buffer (1 M KCl, 5% glycerol, 10 mM imidazole, 50 mM Tris-HCl, 250 mM L-arginine, 1 mM PMSF, pH 7.4) followed by 10 CV wash buffer (binding buffer with 58 mM imidazole). His-tagged protein was then eluted in ∼ 1 CV of elution buffer (binding buffer with 250 mM imidazole), filtered and passed through a 120 mL HiLoad Superdex 200 16/600 pg size exclusion column (Cytiva) pre-equilibrated in storage buffer (0.5 M KCl, 5% glycerol, 20 mM HEPES, 1 mM DTT, pH 7.4). Storage buffer was supplemented with 0.5 mM EDTA for SFPQ ΔN-LCR to prevent degradation by proteolysis from the C-terminus (observed during a preliminary purification attempt). Peak fractions containing H6-mEGFP-tagged protein were pooled, concentrated using a 100 kDa MWCO centrifugal filter (Amicon), flash frozen in 20-50 μL aliquots in liquid nitrogen and stored at - 80°C. To remove the H6-mEGFP tag, tagged protein was incubated with tobacco etch virus protease (TEV; produced in-house) at a 5:1 mass ratio of tagged protein:TEV for 2 hrs at ambient temperature in storage buffer. Untagged SFPQ was then isolated from TEV and H6-mEGFP tag by passing the sample through either a 120 mL HiLoad Superdex 200 16/600 pg or a 24 mL Superdex 200 Increase 10/300 GL size exclusion column (depending on the sample size) pre-equilibrated in storage buffer. Peak fractions containing untagged SFPQ were pooled and concentrated using a 30 kDa MWCO centrifugal filter (Amicon) to a final protein concentration of: 7.9 mg/mL for SFPQ full-length, 0.34 mg/mL for SFPQ ΔN-LCR, 20.5 mg/mL for SFPQ ΔC-LCR, 42.7 mg/mL for SFPQ ΔN&ΔC. Protein concentrations were calculated by absorbance at 280nm using the theoretical extinction coefficient for each protein as calculated from its amino acid sequence using *ProtParam* (Gasteiger *et al*, 2005). Protein purity was assessed by SDS-PAGE (**Figure 2d**).

**Figure 2.**
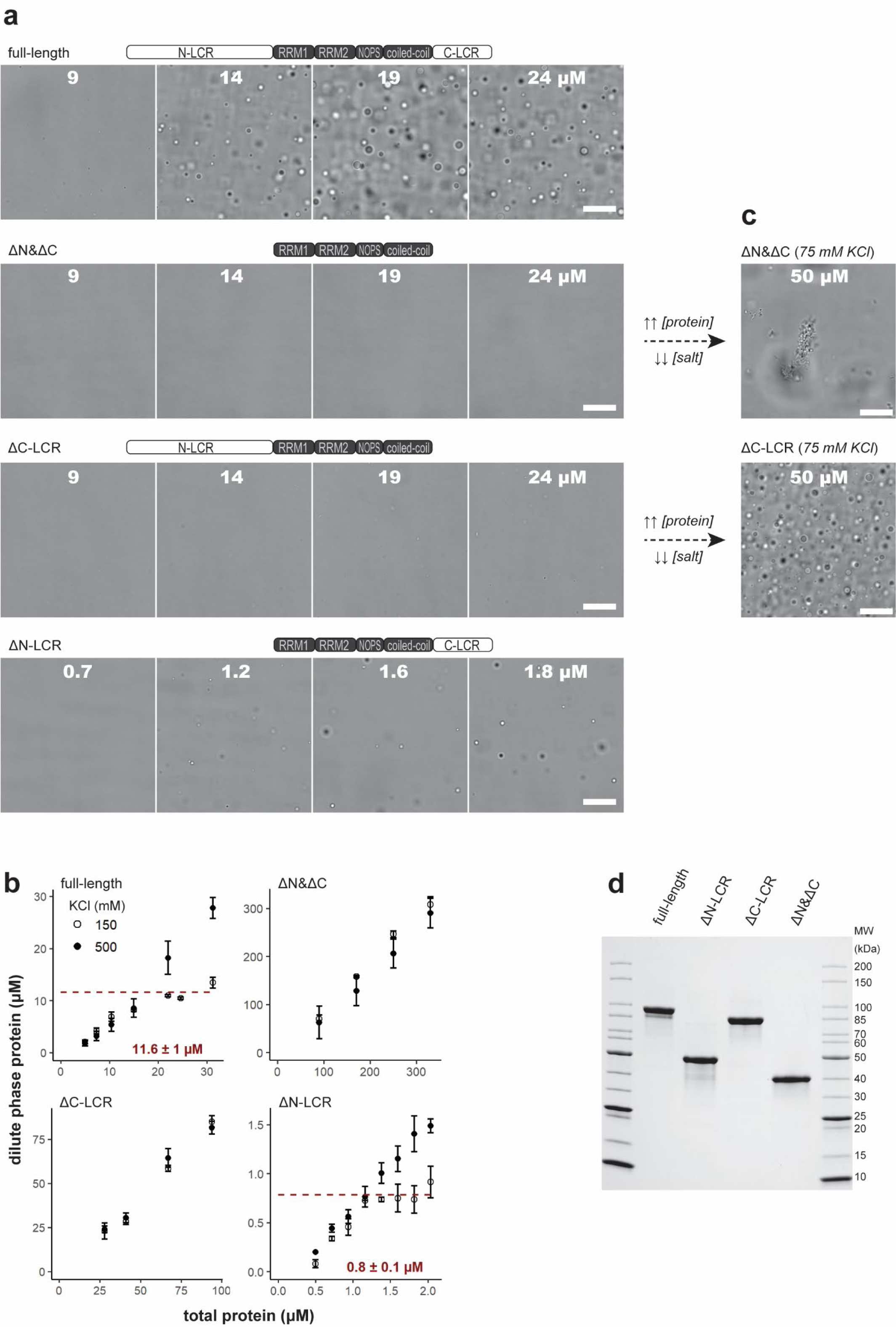
*In vitro* liquid demixing of SFPQ is facilitated by its LCRs. However, the shorter C-LCR is the main driver of condensation, whereas the longer prion-like N-LCR attenuates condensation of the full-length protein. **(a)** Microscopy images show that, at physiological pH and salt concentration (150 mM KCl), both full-length SFPQ (top) and SFPQ ΔN-LCR (bottom) condense to form droplets in a concentration-dependent manner. SFPQ ΔN-LCR condenses at a much lower protein concentration than full-length SFPQ. Removal of both LCRs, or the C-LCR only, (middle panels) abolished droplet formation under these conditions. **(b)** By measuring protein concentration in the dilute phase after dilution to 150 mM KCl, the *C*_sat_ was determined to be 11.6 μM for full-length SFPQ and 0.8 μM for SFPQ ΔN-LCR. For SFPQ ΔC-LCR and ΔN&ΔC, all protein remained in the dilute phase, even at protein concentrations 8- and 28-fold greater than the *C*_sat_ of full-length SFPQ, respectively. Dilutions in protein storage buffer (500 mM KCl) served as a negative control (solid circles). Data points were collected in triplicate; error bars show standard deviation; *C*_sat_ uncertainty expressed as 95% confidence interval. **(c)** Images showing that at high protein concentrations in a solution containing 75 mM KCl, SFPQ ΔN&ΔC formed solid-like fibrous aggregates, whereas SFPQ ΔC-LCR condensed to form spherical liquid droplets. For all microscopy images, the total protein concentration (μM) in each sample is indicated on the image. All scale bars are 20 μm. **(d)** SDS-PAGE of recombinant full-length SFPQ and truncations after purification from bacterial cell culture (Coomassie-stained).

### *In vitro* phase separation assays

Storage buffer was added to protein stocks to produce a series of samples spanning a range of protein concentrations in PCR tubes. To induce phase separation, each sample was then diluted 3/10 by adding low salt buffer (20 mM HEPES, 1 mM DTT, pH 7.4) to give a final solution volume of 10 μL containing 150 mM KCl, 1.5% glycerol, 20 mM HEPES, 1 mM DTT, and a protein concentration of 30% of the pre-diluted sample. Low salt buffer was added rapidly but carefully to avoid bubbles and samples were flick mixed immediately after dilution. For negative controls, storage buffer was added to samples instead of low salt buffer.

To visualise condensed phase droplets in suspension, each sample was transferred to a well of a 384-well non-binding microplate (µClear^®^, Greiner Bio-One) immediately after dilution with low salt buffer and imaged using a Nikon Ti2 inverted microscope with a 60× oil immersion objective. All images were taken with the focal plane ∼ 100 μm above the surface of the bottom of the well. Initially, samples containing mEGFP-SFPQ (full-length) droplets were imaged at 5 min intervals to assess the time-dependence of droplet formation and settling (**Figure S2**). After 10-15 min, droplet size and number appeared to stabilise. After ∼ 35 min, the number of droplets visibly decreased (due to settling). Therefore, all samples were imaged ∼ 20 min post-dilution. For SFPQ ΔN-LCR, initial experiments produced droplets that were very small (∼ 1 μm or less in diameter) making it difficult to visualise droplet shape or fusion. Therefore, samples were rotated slowly for 1 h immediately after transfer to the microplate to allow time for droplets to coalesce in suspension, before imaging as described above.

To determine *C*_sat_ from the dilute phase protein concentration, samples were incubated at room temperature for 10 min after inducing phase separation in PCR tubes as described above. Tubes were centrifuged at 2,000 × *g* for 10 min to accelerate settling of the condensed phase to the bottom of the tube. The protein concentration in the dilute phase was then determined by carefully transferring 2 μL of supernatant (i.e. the dilute phase) to a NanoDrop™ Lite spectrophotometer (Thermo Scientific) and measuring the absorbance at 280nm.

To determine *C*_sat_ from the amount of condensed phase (**Figure S1**), samples containing mEGFP-tagged SFPQ were diluted to 150 mM KCl as described above, transferred immediately to a 384-well microplate, sealed and left overnight on the bench to allow all droplets to settle to the bottom of the plate. Fluorescent images of the surface of the plate were captured using a Nikon Ti2 inverted microscope with a 20× objective and 470nm excitation LED.

Analyses of all microscopy images were performed using ImageJ (v1.53q, NIH). To estimate the relative amount of condensed GFP-tagged SFPQ in samples shown in **Figure S1**, the threshold intensity above which pixels were counted as part of the condensed phase was set using a similar approach to (Wang *et al*., 2018). Briefly, a Gaussian curve was fit to the background peak in the histogram for each image to estimate the mean and standard deviation of the background intensity.

The threshold was then set as the mean background intensity plus three standard deviations. In this way, images could be thresholded in an unbiased way, even if no droplets were visible in the sample. Given that the condensed protein appeared to form puddles on the plate surface (rather than remain as spherical droplets of varying height), the proportion of plate surface area covered by condensed protein was used as a proxy for the relative condensed phase volume.

### Paraspeckle quantitation via fluorescence *in situ* hybridisation

HeLa cells were maintained in DMEM containing high glucose and pyruvate (GIBCO) supplemented with 10% FBS (Sigma Aldrich) and Penicillin-Streptomycin (GIBCO). HeLa cells (7 × 104 cells/well) were seeded onto 1.5 thickness coverslips in a 12-well tray. The following day transfections were performed using 200 ng pEGFP-SFPQ (either full-length, ΔN-LCR, ΔC-LCR or ΔN&ΔC) plasmid, 1 µl P3000 Reagent and 2.5 µl of Lipofectamine 3000 (Thermo Fisher) per well. Transfection media was replaced with fresh media after 6 h. 24 h post-transfection, coverslips were fixed in 4% formaldehyde for 10 min, washed three times with PBS and then permeabilised in 70% ethanol overnight at 4°C. RNA FISH was performed against NEAT1 using NEAT1 Middle Segment with Quasar® 570 Dye (Stellaris FISH probes). After removing 70% ethanol, coverslips were incubated in 1 mL of wash buffer A (Biosearch Technologies) for 5 min. Coverslips were then placed cell side down on 50 µl of hybridization buffer containing probe in a humidified chamber and incubated in the dark at 37°C for 16 h. After hybridisation, coverslips were transferred back to a 12-well tray, cell side up, and washed twice in 1 mL of wash buffer A incubated at 37°C in the dark for 30 min. Coverslips were incubated in buffer B (Biosearch Technologies) containing DAPI for 5 min at room temperature, washed once with buffer B, then mounted onto a glass slide using Vectashield antifade mounting medium (Vector Laboratories) and sealed using nail polish.

Fluorescent images were acquired using a Delta Vision Elite Deconvolution Microscope with a 60× objective. Images were acquired as 70 step z-stacks at intervals of 0.2 μm, with 3D deconvolution and maximum intensity projection performed to generate a final image for analysis. Brightness and contrast of all images were normalised using the ImageJ FIJI software package (NIH). Paraspeckle analysis was performed in CellProfiler using a custom pipeline. In brief, nuclear staining was mapped to determine regions of interest. NEAT1 FISH signal detected within the regions of interest was then used to determine paraspeckle number and area per nuclei. Statistical analysis of paraspeckle number and area was performed using an ordinary one-way ANOVA with multiple comparisons in GraphPad Prism. Line scans to determine colocalisation of NEAT1 and GFP-SFPQ (either full-length, ΔN-LCR, ΔC-LCR or ΔN&ΔC) were performed using the line scan function in ImageJ.

### Western blot

HeLa cells (1.6 × 10^5^ cells/well) were seeded into a 6-well tray. The following day transfections were performed using 500 ng pEGFP-SFPQ (either full-length, ΔN-LCR, ΔC-LCR or ΔN&ΔC) plasmid, 2.5 µl P3000 Reagent and 6 µl of Lipofectamine 3000 (Thermo Fisher) per well. Transfection media was replaced with fresh media after 6 h. 24 h post-transfection cells were washed with cold PBS, detached using a cell scraper, transferred to an Eppendorf tube and centrifuged at 5000 × g for 1 min. The cell pellet was lysed by resuspending in 200 µl RIPA lysis buffer (150 mM NaCl, 25 mM Tris pH 7.5, 1% sodium deoxycholate, 0.1% SDS, 1% IGEPAL CA-630) and incubating on ice for 40 min, with vortexing every 10 min. Samples were centrifuged at 14,000 × g for 15 min, and the supernatant was transferred to another tube. Bradford protein estimation (Bio-Rad #5000006) was performed to ensure equal amounts of protein were loaded for each sample. Protein samples were mixed with SDS gel-loading buffer and heated to 95°C for 10 min. Samples and Precision Plus Protein™ All Blue Prestained Protein Standards (Bio-Rad, 1610373) were loaded onto Mini-PROTEAN TGX Pre-Cast Gels (Bio-Rad, 4561086). Gels were run in Tris/Glycine Buffer (Bio-Rad, 1610771) at 200V. Trans-Blot Turbo Mini Nitrocellulose Transfer packs (Bio-Rad, 1704158) were used for membrane transfer. Membrane was blocked in 5% skim milk PBST. Blocking and antibody incubations were performed for 1 h at room temperature with gentle rocking. GFP antibody (Roche #11814460001) was diluted 1:2,000 in 5% milk PBST, while secondary (horseradish-peroxidase conjugated) goat anti-mouse IgG H&L HRP (Abcam, ab97023) was diluted 1:10,000 in 5% milk PBST. Three PBST washes with gentle rocking were conducted for 10 min each after each antibody incubation. Luminata Crescedo Western HRP substrate (Merck, WBLUR0100) was added and blot images were acquired by Bio-Rad Chemidoc. Bio-Rad Imagelab Version 6.1 was used to quantify total protein levels and the intensity of the protein chemiluminescent bands. The relative intensity of each chemiluminescent band was normalized to the amount of total protein in the sample lane, and the sizes of chemiluminescent bands determined in relation to the standards ladder bands.

### DNA damage site recruitment assay

HeLa cells were transfected with pEGFP-SFPQ constructs (full-length, ΔN-LCR, ΔC-LCR or ΔN&ΔC) for 48 h. A day prior to the experiment, cells were treated with 10 μM BrdU in DMEM without phenol red (Gibco) containing 10% FBS and 1% Pen/Strep. The recruitment of GFP-SFPQ was imaged at 5 Hz on a Zeiss LSM710 confocal microscope. DNA damage was induced by irradiating 10 pixel-width of the nucleus with a 405 nm laser (line scan mode). The laser was set with a line attenuator transmission of 92%, 1 iteration and a pixel dwell time of 177.32 µs. Regions of interest were drawn to the DNA damage site to measure the GFP fluorescence over time using the ImageJ software (NIH). DNA damage-induced changes in fluorescence were calculated by normalizing the background subtracted values to the baseline fluorescence prior to laser microirradiation (ΔF/F_o_).

### *In silico* sequence analysis

The *IUPred2A* webserver (https://iupred2a.elte.hu/) (Erdős & Dosztányi, 2020) was used with default parameters (IUPred2 long disorder) to predict intrinsically disordered regions within the amino acid sequence of human SFPQ (UniProt accession P23246). The *PLAAC* webserver (http://plaac.wi.mit.edu/) (Lancaster *et al*., 2014) was used to find prion-like regions, with a minimal length (‘core length’, *c*) of 60 residues (default). The α value was set to 0.5, such that the background amino acid frequencies used were the average of the frequencies across the entire *Homo sapiens* and *Saccharomyces cerevisiae* proteomes, as recommended by the program developers when scanning non-*S. cerevisiae* proteins (Lancaster *et al*., 2014).

## RESULTS

### SFPQ can form a condensed liquid-like phase *in vitro*

The ability to form homotypic liquid-like droplets *in vitro* has been described previously for many RNA-binding proteins with extended LCRs (Maharana *et al*, 2018; Wang *et al*., 2018) (see also (Boeynaems *et al*, 2018; Borcherds *et al*., 2021) for reviews). We therefore first sought to determine the phase behaviour of SFPQ *in vitro*. Recombinant full-length human SFPQ with an N-terminal GFP tag was purified from bacterial cell culture; the GFP tag was removed and untagged SFPQ was concentrated in a buffer containing 500 mM KCl (**Figure 2d**). Upon dilution to physiological salt concentration (150 mM KCl), micron-sized droplets formed in a concentration-dependent manner (**Figure 2a**). As droplets settled, they were observed to fuse with the bulk condensed phase at the bottom of the sample container, confirming their liquid-like state (**Movie S1**). Furthermore, by measuring the absorbance at 280 nm of the supernatant after centrifuging to remove all droplets, we could determine the protein concentration in the dilute phase (i.e. the protein concentration outside of droplets). We observed that the dilute phase protein concentration was unchanged in samples with total protein concentrations greater than the concentration at which droplets began to form (**Figure 2b**). This behaviour is consistent with the expectation for phase separation in a binary mixture. We identify the invariant concentration measured in the coexisting dilute phase with the saturation concentration (*C*_sat_) (Alberti *et al*, 2019). Under these conditions (150 mM KCl, pH 7.4, ambient temperature), we determined a *C*_sat_ of 11.6 μM for full-length SFPQ. This *C*_sat_ is in the same range as the estimated nuclear concentration of similar RNA-binding proteins (FUS, EWSR1, TAF15, TDP43 hnRNPA1 (Maharana *et al*., 2018)). The total concentration of SFPQ in HeLa cells has been estimated using mass spectrometry to be 1.8 μM (Hein *et al*, 2015). However, given that SFPQ is predominantly localised to the nucleus, its concentration within the nucleus is likely to be ∼ ten-fold greater than this. Furthermore, the increase in local concentration due to recruitment to DNA and/or RNA sites, along with the highly crowded intracellular environment, makes condensation of SFPQ in the nucleus very plausible.

The method described above for *C*_sat_ determination relies on measuring the protein concentration in the dilute phase. To further verify this result, we used an alternative method to instead quantify the amount of condensed phase. For this, we used GFP-tagged SFPQ to facilitate imaging by fluorescence. By imaging after all droplets had settled to the bottom of the sample, we were able to quantify the relative amount of condensed phase in each sample over a range of concentrations (**Figure S1a** and **S1b**). For samples containing droplets, the relationship between amount of condensed phase and total protein concentration was approximately linear, as expected for a binary two-phase system (Alberti *et al*., 2019). The minimum concentration required for phase separation to occur (i.e. the *C*_sat_) was then calculated by extrapolation to zero. For comparison we also determined the *C*_sat_ of GFP-SFPQ using the centrifuge-based assay as described above (**Figure S1c**). Both methods gave a *C*_sat_ for GFP-SFPQ of 1.5-1.6 μM, providing confidence in the accuracy of these approaches. This also showed that the N-terminal GFP tag increases the propensity of SFPQ to condense *in vitro*. Therefore, for further *in vitro* experiments we used untagged proteins only and the centrifuge-based assay for *C*_sat_ determination.

### The C-terminal LCR of SFPQ drives phase separation, which is attenuated by the N-terminal prion-like LCR *in vitro*

To determine the contribution of each LCR of SFPQ independently on the *in vitro* condensation behaviour of the protein, recombinant SFPQ truncations lacking the N-LCR, C-LCR or both N- and C-LCR were purified from bacterial cell culture (**Figure 2d**) and assayed for *in vitro* phase separation in the same way as for full-length SFPQ. Unsurprisingly, deletion of both N- and C-LCRs (SFPQ ΔN&ΔC) produced a protein that would not phase separate to form droplets (**Figure 2a**), remaining as a single soluble phase even at a protein concentration almost 30-fold greater that the *C*_sat_ of full-length SFPQ (**Figure 2b**). Even after further dilution to lower the KCl concentration to 75 mM, SFPQ ΔN&ΔC remained soluble up to a concentration of ∼ 50 μM, at which point it precipitated as visible solid-looking aggregates (**Figure 2c**), indicating that without its LCRs, SFPQ cannot form a condensed liquid phase under physiological-like conditions (ionic strength, pH, [protein], temperature) *in vitro*.

Deletion of the C-LCR only (amino acids 599-707) also resulted in a protein which would not phase separate under the same conditions as for the full-length protein (**Figure 2a**). Even at a total protein concentration eight-fold greater than the *C*_sat_ of full-length SFPQ, SFPQ ΔC-LCR remained as a single dilute phase, as determined using the centrifuge assay (**Figure 2b**). This clearly implies that the C-LCR is crucial for driving SFPQ phase separation. To assess whether SFPQ ΔC-LCR could phase separate at all, we decreased the salt concentration further by dilution. Liquid droplets were clearly observed to form in a sample containing 75 mM KCl and 50 μM SFPQ ΔC-LCR (**Figure 2c** and **Movie S2**). This is in contrast to SFPQ ΔN&ΔC, which precipitated as a solid under these conditions, demonstrating that the N-LCR of SFPQ can facilitate condensation of the protein as a liquid, albeit requiring more extreme conditions (i.e. less like physiological ionic strength and [protein]) than for full-length SFPQ.

Unexpectedly, deletion of the longer prion-like N-LCR (amino acids 1-275) resulted in a protein with an increased propensity to phase separate, suggesting that the N-LCR attenuates demixing (promotes mixing) of SFPQ in the context of the full-length protein. The *C*_sat_ of SFPQ ΔN-LCR in 150 mM KCl buffer was determined to be 0.8 μM (**Figure 2b, bottom-right**); a ∼15-fold decrease as compared to the full-length protein. Consistent with an increased propensity to self-associate, we observed turbidity while concentrating SFPQ ΔN-LCR in storage buffer containing 500 mM KCl, and therefore, could not access protein concentrations comparable to the full-length protein (note the lower concentration range for ΔN-LCR in **Figure 2a** and **2b**). The SFPQ ΔN-LCR droplets that formed upon dilution to 150 mM KCl were smaller and more sparse than for the full-length protein (**Figure 2a, bottom**), as expected for a lower condensed volume in samples containing protein concentrations that we could readily achieve. Despite this, droplet fusion events could still be observed, indicating that this condensed phase formed by SFPQ ΔN-LCR was liquid-like (**Movie S3**).

Together, these *in vitro* data suggest that the LCRs of SFPQ have distinct roles in determining the phase behaviour of the protein. Independently, the N-LCR and the C-LCR can each facilitate homotypic condensation of the conserved globular DBHS region of SFPQ as a liquid; in the absence of both LCRs, the protein precipitates at high concentrations as a solid. However, the shorter C-LCR is clearly the main driver of condensation, whereas, in the absence of the C-LCR, much more extreme conditions are required to observe liquid condensation mediated by the longer prion-like N-LCR. Perhaps most significantly, we find that the N-LCR in fact attenuates SFPQ condensation in the context of the full-length protein at physiological salt concentration.

### The C-terminal LCR of SFPQ is critical for paraspeckle formation *in vivo*, whereas the N-LCR increases SFPQ nuclear dispersal

It has been observed that prion-like domains are enriched in human nuclear DNA- and RNA-binding proteins (March *et al*., 2016). Prion-like regions have been suggested to be important mediators of functional aggregation to form biomolecular condensates, with proteins containing prion-like LCRs shown to be over-represented in paraspeckles (Hennig *et al*., 2015) – membraneless organelles present in the nucleus of many cell types. Paraspeckles can be either singular spheroids of 360 nm diameter, or chains of spheroids with a constant width, and length up to 1.5 μm. SFPQ is one of seven proteins shown to be indispensable for the formation of paraspeckles (Naganuma *et al*, 2012). We have previously used FISH to specifically locate the long non-coding RNA, NEAT1_2, upon which paraspeckles are scaffolded (Lee *et al*., 2015; Lee *et al*., 2022; West *et al*., 2016). In addition, SFPQ subnuclear localisation has been observed in live cells previously by over-expression of GFP-tagged SFPQ (Dye & Patton, 2001). Therefore, to probe for roles of the LCRs of SFPQ in subnuclear organisation *in vivo*, we performed quantitative analysis on paraspeckles – identified as NEAT1 FISH puncta – in cultured HeLa cells overexpressing GFP-tagged SFPQ truncations (**Figure 3**). Due to the nature of transient transfection, expression of each protein was variable when comparing individual cells. However, the localisation patterns of each protein within the nucleus (discussed below) were qualitatively the same in cells with low, medium, or high exogenous expression (**Figure S3**). Importantly, Western blot analysis showed that transient transfection resulted in comparable average expression levels for all four proteins in each cell culture (**Figure S4**), allowing us to directly compare paraspeckle quantitation data for each cell population.

**Figure 3.**
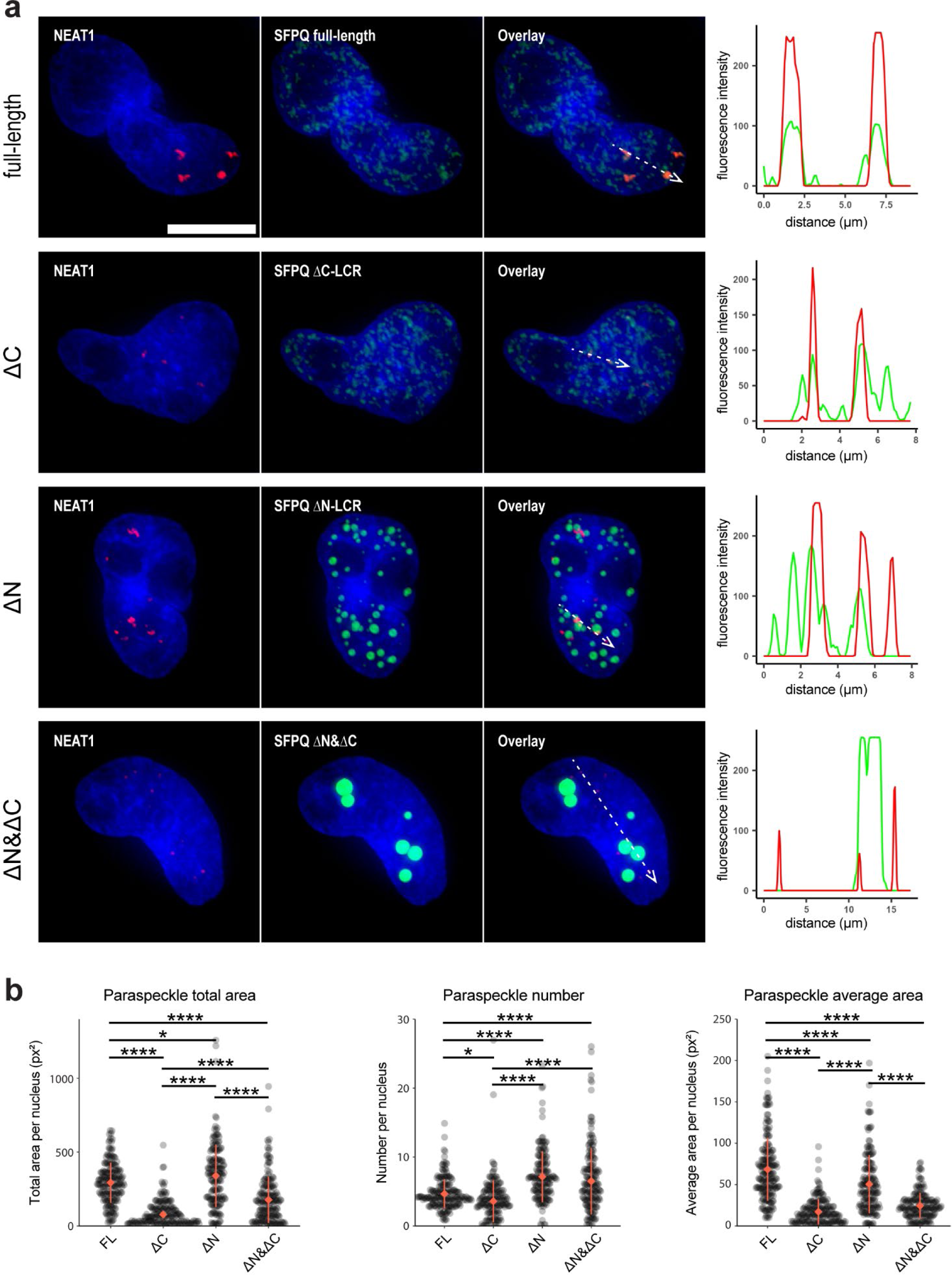
Contrasting effects of over-expressing SFPQ LCR truncations on nuclear paraspeckle size and abundance. **(a)** Representative fluorescence images of Hela cells overexpressing GFP-tagged full-length SFPQ, ΔC-LCR, ΔN-LCR and ΔN&ΔC. NEAT1_2 FISH (red) shows the location of paraspeckles. Intensity profiles (dashed arrows) highlight the localisation of GFP-tagged protein relative to paraspeckles (see Figure S5 for additional examples). SFPQ ΔC-LCR has a similar nuclear localisation pattern to full-length. Removal of the N-LCR results in a large number of droplet-like non-paraspeckle puncta. Scale bar = 10 μm. **(b)** Quantification of paraspeckle size and number in cells expressing GFP-tagged SFPQ truncations shows that removing the C-LCR results in fewer and smaller paraspeckles, whereas ΔN-LCR results in a modest increase in total paraspeckle area compared to full-length. Expression of SFPQ ΔN&ΔC also reduces paraspeckle area, but not to the same extent as ΔC-LCR (**** p<0.0001, and * p<0.05 by ANOVA).

SFPQ is a multifunctional nuclear protein, and localises along with other essential paraspeckle proteins, NONO and FUS, within the core of paraspeckles in cells where they are present (Yamazaki *et al*, 2018). Therefore, as expected, in cells expressing full-length SFPQ, GFP signal was distributed throughout the nucleus, and co-localised with NEAT1_2 where NEAT1_2 was present (**Figure 3a**; see **Figure S5** for additional linescans). To determine the effect of SFPQ over-expression on paraspeckle number and size, we quantitated total paraspeckle area (i.e. the sum of pixels with NEAT1_2 FISH signal), number of individual paraspeckles, and the average area of individual paraspeckles (total paraspeckle area/paraspeckle number) per nucleus in cells over-expressing either GFP, or GFP-SFPQ. Compared to GFP alone, cells over-expressing GFP-SFPQ had a small but significant decrease in paraspeckle number, accompanied by an increase in paraspeckle area by approximately 1.5 to 2-fold (**Figure S6**). Thus, over-expressed GFP-SFPQ is targeted to paraspeckles and increases their average size but does not increase singular paraspeckle numbers.

To determine the effect of removing the C-LCR from SFPQ on paraspeckle abundance, a truncation was constructed lacking amino acids 599-699; the C-terminal NLS (amino acids 700-707) was included to ensure nuclear localisation. GFP signal was distributed throughout the nucleus, and co-localised with NEAT1_2, similar to the full-length protein (**Figure 3a**). However, compared to cells expressing full-length SFPQ, cells expressing SFPQ ΔC-LCR showed a marked decrease in paraspeckle area, accompanied by a small but significant decrease in number of paraspeckles (**Figure 3b**). Given the presence of endogenous DBHS proteins (i.e. no knock-down was performed), this demonstrates that SFPQ ΔC-LCR acts as a dominant negative in regards to paraspeckle size and abundance, consistent with our *in vitro* results showing that the C-LCR drives SFPQ condensation.

In contrast to the C-LCR, which is clearly important for paraspeckle maintenance, loss of the N-LCR (1-275) resulted in a small but significant increase in total paraspeckle area per nucleus; a combination of an increase in the number of paraspeckles, accompanied by a small decrease in their average size (**Figure 3b**). Compared to full-length SFPQ and the ΔC-LCR truncation, SFPQ ΔN-LCR showed a different localisation pattern within the nucleus, with GFP signal isolated to bright puncta, most of which were not co-localised with NEAT1 and therefore not paraspeckles (**Figure 3a**). These non-paraspeckle puncta were of various sizes (∼ 0.2 - 2 µm) and all approximately spherical. This is consistent with our *in vitro* data showing that SFPQ ΔN-LCR has a greater tendency to phase separate to form condensed droplets. Similar bright non-paraspeckle puncta have been observed for SFPQ truncations missing the N-LCR previously (Dye & Patton, 2001; Ha *et al*, 2011). Interestingly, NEAT1 signal did not co-localise with SFPQ ΔN-LCR as consistently as for full-length and ΔC-LCR, suggesting that the N-LCR may be important for localisation of SFPQ to paraspeckles (**Figure 3a** and **Figure S5**).

Finally, over-expression of a double-deletion missing both LCRs (SFPQ ΔN&ΔC; includes NLS) resulted in a decrease in total paraspeckle area compared to full-length SFPQ; a combination of a small increase in number (as for ΔN-LCR), but a large decrease in average paraspeckle area (as for SFPQ ΔC-LCR) (**Figure 3b**). However, the total paraspeckle area and number were significantly greater than for SFPQ ΔC-LCR. Our interpretation from this comparison between SFPQ ΔN&ΔC and SFPQ ΔC-LCR is that inclusion of the N-LCR has an inhibitory effect on paraspeckle formation. In contrast to the wild-type like nuclear localisation observed for SFPQ ΔC-LCR, the double deletion appeared as discrete, round, non-paraspeckle puncta, similar to SFPQ ΔN-LCR (**Figure 3a**). This is again consistent with the N-LCR attenuating SFPQ demixing, increasing its tendency to distribute throughout the nucleus rather than condensing to form discrete puncta. Given that the ability to condense is very likely to be important for driving paraspeckle formation, this suggests that SFPQ ΔC-LCR acts as a stronger dominant negative than SFPQ ΔN&ΔC due to the inclusion of the N-LCR. Similar to SFPQ ΔN-LCR, NEAT1 signal did not consistently co-localise with SFPQ ΔN&ΔC, pointing again to the N-LCR as important for localisation of SFPQ to paraspeckles (**Figure 3a** and **Figure S5**). DBHS proteins have been previously observed to localise to the periphery of the nucleolus (‘perinucleolar caps’) upon inhibition of transcription (Fox *et al*, 2005; Fox *et al*, 2002). Here, nucleoli could be identified as dark regions (less DAPI stain) within the nucleus (**Figure S7**). There was no clear overlap between these regions and GFP, showing that the condensates containing SFPQ ΔN&ΔC are distinct from nucleoli.

### The C-terminal LCR is critical for condensation of SFPQ at DNA damage sites

High avidity liquid-like interactions between proteins with intrinsically disordered regions and nucleic acids have been proposed as a key mechanism driving macromolecular condensation at specific sites in the cell nucleus (Feric *et al*., 2016; Maharana *et al*., 2018; Morin *et al*., 2022). An example is the assembly of DNA repair machinery at DNA damage sites (Levone *et al*, 2021). Previous studies have shown that SFPQ is directly involved in DNA damage repair via the non-homologous end joining (NHEJ) pathway. *In vitro*, the SFPQ-NONO dimer forms a committed pre-ligation complex with another NHEJ factor (Ku) and the DNA substrate, which is essential for DNA ligase IV/XRCC4-mediated joining of double-stranded DNA ends (Bladen *et al*, 2005). *In vivo*, depletion of SFPQ severely delays repair of double-strand breaks (Salton *et al*, 2010), and results in a marked increase in radiation-sensitivity in HeLa cells (Ha *et al*., 2011). In addition, SFPQ can bind to DNA in the absence of other DBHS proteins (Lee *et al*., 2015; Song *et al*, 2005), and is the DBHS partner which drives DNA binding *in vitro* (Zhang *et al*, 1993) and *in vivo* (Ha *et al*., 2011). Therefore, to probe for roles of the LCRs of SFPQ in driving condensation at functional sites in the cell nucleus, we assessed the recruitment of SFPQ to DNA damage sites in HeLa cells overexpressing GFP-tagged SFPQ.

Within minutes of causing localised damage to DNA in the cell nucleus using a laser microbeam, GFP signal accumulated at damage sites, indicating rapid SFPQ recruitment, as shown previously (Ha *et al*., 2011; Salton *et al*., 2010) (**Figure 4a**). GFP signal peaked approximately 3 minutes after inducing DNA damage and decreased gradually back to baseline after 10-15 minutes (**Figure 4b**). Removal of the C-LCR reduced the relative recruitment by approximately two-fold; the magnitude of this effect was similar for both SFPQ ΔC-LCR and SFPQ ΔN&ΔC. Interestingly, although DNA damage recruitment was unchanged for SFPQ ΔN-LCR, additional bright GFP puncta appeared to be present in these cells compared to those expressing full-length SFPQ (**Figure 4a, row 3**), consistent with the additional GFP puncta observed for SFPQ ΔN-LCR in our paraspeckle quantitation assay (**Figure 3a**). Compared to full-length SFPQ, no significant difference in recruitment to damage sites was observed for SFPQ ΔN-LCR (**Figure 4c**). Together, these results suggest that deletion of the C-LCR, but not the N-LCR, impairs SFPQ condensate formation at DNA damage sites. Conversely, removing the N-LCR increases the condensation propensity of SFPQ in the nucleus, consistent with our *in vitro* data.

**Figure 4.**
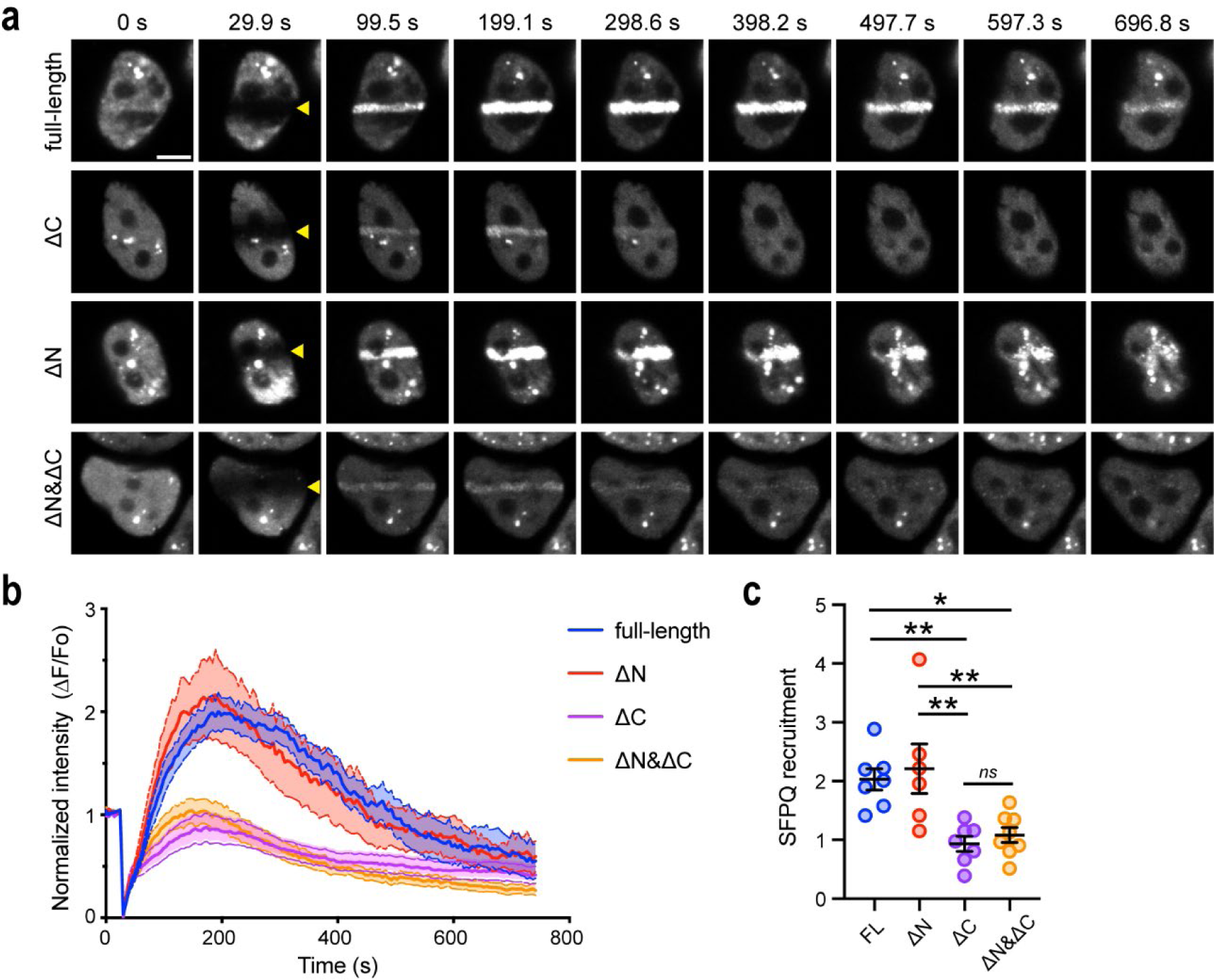
Efficient recruitment of SFPQ to DNA damage sites *in vivo* requires its C-LCR, but not its N-LCR. **(a)** HeLa cells overexpressing GFP-SFPQ, either full-length (FL), ΔN-LCR, ΔC-LCR or ΔN&ΔC-LCR, were subjected to laser microirradiation. Scale bar = 5 μm. **(b)** Traces showing relative changes in GFP fluorescent intensity (ΔF/Fo) at sites of DNA damage over time. Solid and dashed lines indicate means and SEM, respectively. FL, n = 7 cells; ΔN-LCR, n = 6 cells; ΔC-LCR, n = 7 cells; and ΔN&ΔC-LCR, n = 8 cells. **(c)** Peak fluorescent intensity (SFPQ recruitment) at DNA damage sites. Data are presented as means ± SEM. *p<0.05, **p<0.01 (one-way ANOVA with Tukey’s multiple comparison test).

## DISCUSSION

Demixing of proteins and nucleic acids is emerging as a key mechanism describing the dynamic and reversible organisation of molecules in the cell. The central involvement of nucleic acid-binding proteins with disordered, low-complexity regions – particularly those with prion-like composition – tends to be a common feature of many studies to date, with condensate formation driven by prion-like domains emerging as a key theme (Dorone *et al*, 2021; Fox *et al*., 2018; Franzmann *et al*, 2018; Harrison & Shorter, 2017; Hennig *et al*., 2015; Levone *et al*., 2021; West *et al*., 2016). SFPQ is an abundant, ubiquitous and multifunctional RNA- and DNA-binding protein with an extended N-terminal LCR – approximately one quarter of which is classified as prion-like. Unexpectedly, here we show that rather than driving phase separation, this prion-like N-LCR attenuates SFPQ phase separation, which is in fact driven by the shorter, non-prion-like, C-terminal LCR. We have observed this using *in vitro* droplet-forming assays and using two different cell-based assays quantifying two different functional biomolecular condensates: paraspeckles and DNA damage repair foci.

Our *in vitro* data show that, under the range of conditions tested herein, the core globular DBHS region of SFPQ alone cannot condense as a liquid, and instead precipitates at high concentrations as solid-like aggregates. Under the same conditions, the addition of the prion-like N-LCR allows the protein to condense as a liquid instead. However, our results urge caution regarding the interpretation of observed *in vitro* protein liquid condensation: in the absence of the C-LCR, much more extreme conditions (i.e. higher protein concentration and lower ionic strength) were required to observe droplet formation than for the full-length protein, indicating a large decrease in the propensity to undergo demixing compared to the full-length protein. That is, although the N-LCR can *allow* condensation of the protein as a liquid under conditions in which the protein would otherwise precipitate as a solid, it is not the *driver* of protein condensation under physiological conditions. Indeed, even though our SFPQ ΔC-LCR truncation contained all of the domains thus far identified as essential for DNA-binding, RNA-binding and paraspeckle localisation (Knott *et al*., 2022; Lee *et al*., 2015; Wang *et al*, 2022), we observed significant impairment in both DNA damage site localisation and paraspeckle formation, indicating that alone the N-LCR is insufficient to drive functional condensation of the protein in the cell. The C-LCR also allowed the protein to condense as a liquid *in vitro*. However, in contrast to the N-LCR, the C-LCR caused the core DBHS region to condense at much lower protein concentrations, under conditions much closer to physiological. Furthermore, to our surprise, we found that removing the prion-like N-LCR from the full-length protein resulted in a significant increase in the phase separation propensity *in vitro*, and the formation of additional condensates *in vivo*. Together, this shows that the C-LCR drives the formation of functional condensates, whereas the N-LCR attenuates condensation of the full-length protein. This is, to our best knowledge, the first description of an extensive prion-like LCR that serves to regulate rather than drive a protein’s condensation propensity; most recent studies on LCRs of RNA-binding proteins have focused solely on their roles in promoting protein phase separation.

It is well established that DBHS proteins are obligate dimers – able to form homo- and heterodimers – with a preference for heterodimerisation (Huang *et al*., 2018; Knott *et al*., 2022; Laurenzi *et al*, 2022; Lee *et al*., 2022). SFPQ, NONO and PSPC1 co-immunoprecipitate (Bottini *et al*, 2017; De Silva *et al*, 2019), and SFPQ is essential for the localisation of NONO both to DNA damage sites (Ha *et al*., 2011) and to paraspeckles (Lee *et al*., 2022). Previous in-depth structural studies have identified the minimal region necessary and sufficient for DBHS dimer formation as composed of RRM2, NOPS and coiled-coil domains (Lee *et al*., 2015; Passon *et al*., 2012). It is important to note that the entire core conserved DBHS region, including all domains necessary for dimer formation, was left intact in the truncations of SFPQ used herein. This provides a basis for interpreting the results of our *in vivo* experiments, in which SFPQ truncations were over-expressed in the presence of endogenous wild-type DBHS proteins and therefore would be very likely to exist as heterodimers with endogenous DBHS proteins in the cell.

The putative DNA binding domain (DBD) of SFPQ, which is part of the N-LCR and loosely defined as amino acids 214-275, has been shown to be essential for DNA binding *in vitro* (Lee *et al*., 2015) and localisation of SFPQ/NONO and other NHEJ factors to DNA damage sites *in vivo* (Ha *et al*., 2011). This latter study used cells in which endogenous SFPQ was knocked-down (Ha *et al*., 2011). In contrast, our results show that SFPQ ΔN-LCR, for which most of the DBD is absent, localises to DNA damage sites to the same degree as full-length SFPQ. This suggests that interactions with endogenous SFPQ can compensate for the absent DBD of SFPQ ΔN-LCR, allowing for the wild-type-like DNA damage localisation we observe. In addition to direct SFPQ-DNA interactions, it is likely that other factors also contribute to SFPQ recruitment to DNA damage sites: Recruitment of the RNA-binding protein FUS to DNA damage sites precedes SFPQ, and its absence strongly reduces SFPQ recruitment (Levone *et al*., 2021). In light of this, the significant impairment of DNA-damage localisation we observe upon deletion of the C-LCR is striking, as it shows that even the presence of endogenous SFPQ cannot compensate for the lack of this domain. This suggests that DBHS dimers containing SFPQ ΔC-LCR have a lower propensity to condense than those that contain full-length SFPQ. That is, SFPQ ΔC-LCR attenuates condensation, even in the presence of endogenous proteins. This is consistent with our paraspeckle quantitation data showing that over-expressed SFPQ ΔC-LCR impairs paraspeckle formation. Furthermore, the fact that SFPQ ΔN&ΔC localised to discrete rounded puncta, rather than distributed throughout the nucleus like SFPQ ΔC-LCR, points again to the N-LCR as a region that attenuates condensation of DBHS proteins in the cell.

Altogether, this points to a regulatory role for the N-LCR, rather than a role as a driver of phase separation. Interestingly, data from two independent studies involving SFPQ truncations missing the N-LCR have hinted at this previously. In a study focusing on nuclear speckles, upon over-expression of SFPQ truncations lacking amino acids 1-278 or 1-337 in Hela cells, the authors observed discrete nuclear puncta, which appeared larger and more rounded than nuclear speckles (Dye & Patton, 2001). Similarly, authors of a study focussing on the localisation of SFPQ to DNA damage sites noted the accumulation of SFPQ truncations lacking 1-297 or 1-372 in discrete, bright non-paraspeckle puncta (Ha *et al*., 2011). These observations are consistent with our results and suggest that, rather than driving condensation, the prion-like N-LCR is required to prevent aberrant protein condensation in the nucleus.

Despite being much shorter than the N-LCR, we show that the C-LCR is the main driver of homotypic liquid demixing *in vitro*. This same region is critical for both recruitment to DNA damage sites and paraspeckle formation *in vivo*, strongly suggesting that liquid phase condensation of SFPQ, via its C-LCR, is essential for achieving its normal functions in the cell nucleus. Further studies aiming to dissect the sequence features within the C-LCR important for driving demixing are currently underway. In addition, it is expected that interactions with nucleic acids via the DNA binding and RRM domains of SFPQ, and also polymerisation via the coiled-coil (Lee *et al*., 2015), will emerge as important contributors to the condensation of DBHS proteins with other macromolecules within the complex intracellular environment.

In summary, our data provide nuance to the emerging paradigm regarding phase separation driven by low-complexity prion-like domains by showing that the two LCRs of SFPQ have contrasting and possibly opposing effects on condensate formation. It is striking that the shorter C-LCR appears to be the main driver of condensation, whereas the much longer N-LCR of SFPQ, which makes up about one-third of the total polypeptide length and contains a long prion-like region, attenuates condensation of the full-length protein. It may be plausible to propose a model in which direct interaction between the different LCRs of SFPQ are dynamically modulated to provide a mechanism for spatial and temporal regulation of condensation (**Figure 5**). That is, in the absence of competing factors, the long N-LCR attenuates SFPQ condensation by directly interfering with intermolecular interactions mediated by the C-LCR. Conversely, in the presence of competing factors such as nucleic acids or other protein interaction partners which occupy the N-LCR, the C-LCR is then free to form intermolecular interactions to drive demixing leading to condensate formation. This presents a mechanism in which phase separation can be regulated by competing intra- and intermolecular interactions between different LCRs within the same protein. A conceptually similar mechanism has been proposed recently for the self-regulation of phase separation of the stress-granule protein, G3BP1 (Guillén-Boixet *et al*, 2020). The highly variable N- and C-terminal LCRs of the three human DBHS homologs, coupled with their intrinsic combinatorial heterodimerisation nature, present additional avenues for complex self-regulation of condensate formation in the cell. This, along with their multifunctional nature, present DBHS proteins as an attractive model system for describing relationships between LCR composition and the emergent material state of proteins that contain intrinsically disordered regions.

**Figure 5.**
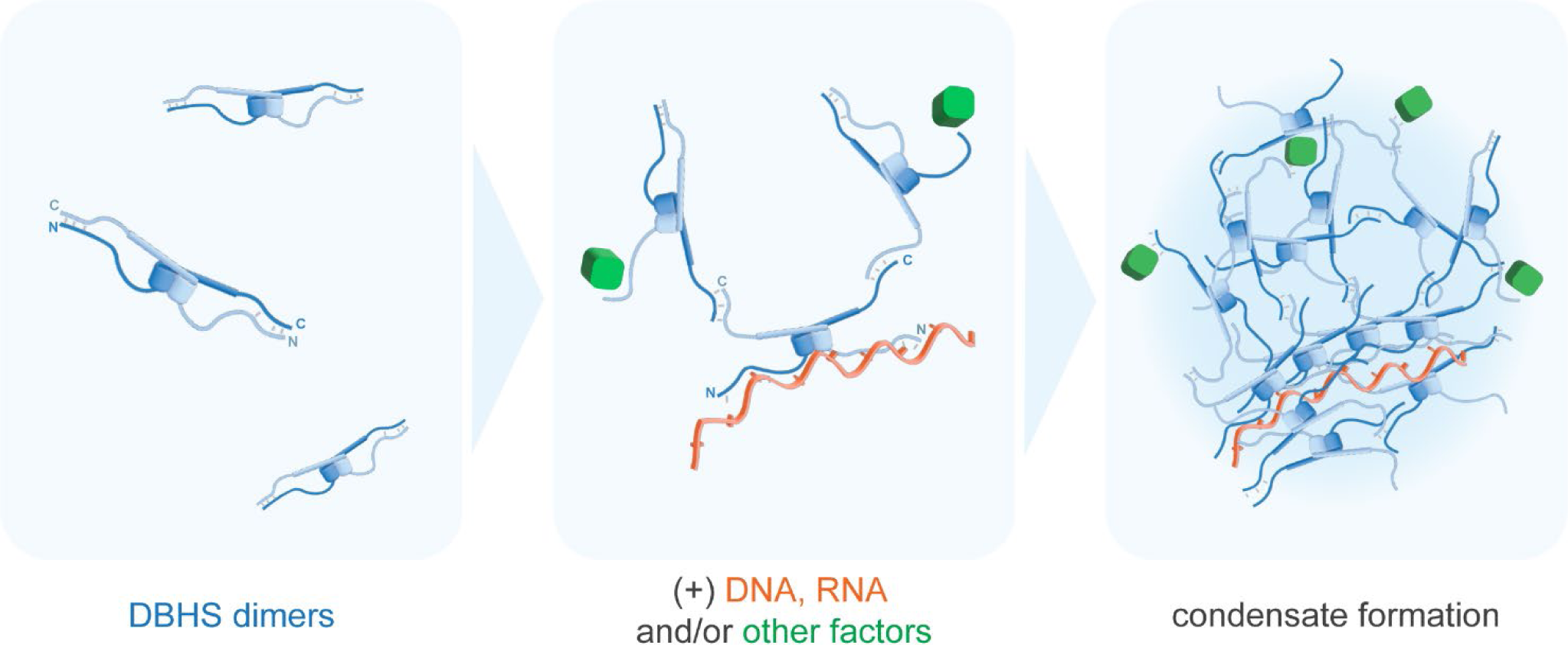
Self-regulation of DBHS protein phase separation via competing intra- and intermolecular interactions between N- and C-terminal LCRs. In the absence of other factors, direct intra-dimer interactions between the N- and C-LCRs preclude inter-dimer interactions, preventing demixing (left). Interactions with DNA, RNA and/or other factors sequester the N-LCR (middle), allowing the C-LCR to form productive inter-dimer interactions, nucleating phase separation to facilitate condensate formation (right).

## DATA AVAILABILITY

The ImageJ macro used to threshold fluorescent images and calculate condensed area is available from https://github.com/acmarshall88/Drop-Counter/blob/main/Drop_Counter.ijm. Fitting of linear models for calculation of *C*_sat_ from centrifuge assay data and fluorescent image data was performed using simple custom scripts written in R (v4.1.0), available from https://github.com/acmarshall88/Csat-CalculatoRs.

## SUPPLEMENTARY DATA

Supplementary Data are available at NAR Online.

## ACKNOWLEDGEMENTS

The authors thank both Dr Hiroyuki Uechi and Dr Patrick McCall (Max Planck Institute of Molecular Cell Biology and Genetics, Dresden) for helpful feedback while drafting the manuscript. The authors are grateful for free access to UCSF ChimeraX (v1.3) (Pettersen *et al*, 2021) (used for molecular graphics in **Figure 1b**), developed by the Resource for Biocomputing, Visualization, and Informatics at the University of California, San Francisco, with support from NIH R01-GM129325 and the Office of Cyber Infrastructure and Computational Biology, NIAID. Imaging was performed at the Queensland Brain Institute’s Advanced Microscopy Facility, supported by the Australian Government through ARC LIEF grant LE130100078.

## Author contributions

M.L., A.H.F., C.S.B. and A.C.M. designed the project. A.C.M. and J.C. purified proteins and performed *in vitro* experiments with guidance from M.L. S.K. performed HeLa cell culture, imaging and analysis. T.Z., J.W. and V.A. performed DNA damage recruitment assays. A.C.M. drafted the manuscript. All authors contributed to data interpretation, manuscript review and editing.

## SUPPLEMENTARY DATA

**Movie S1. Spherical droplets formed by SFPQ *in vitro* display liquid-like properties.** As droplets settle they fuse with bulk condensed protein at the bottom of the plate. Droplets were observed to fade rapidly from view (arrows) *only* in this focal plane (which is higher than the plate surface), and no droplets were visible below this focal plane, consistent with this being the surface of a bulk condensed liquid phase.

**Movie S2. SFPQ ΔC-LCR droplets fusing in suspension.** Sample contains a total SFPQ ΔC-LCR concentration of 50 μM in buffer containing 75 mM KCl.

**Movie S3. SFPQ ΔN-LCR droplets fusing on the surface of the plate.** Sample contains a total SFPQ ΔN-LCR concentration of 1.8 μM in buffer containing 150 mM KCl.

**Movie S4.** Laser microirradiation induces rapid recruitment of GFP-SFPQ full-length to DNA damage sites in HeLa cells.

**Movie S5.** Laser microirradiation induces rapid recruitment of GFP-SFPQ ΔN-LCR to DNA damage sites in HeLa cells.

**Movie S6.** GFP-SFPQ ΔC-LCR is recruited to DNA damage sites to a significantly lesser extent. SFPQ ΔC-LCR also forms additional spherical condensates.

**Movie S7.** GFP-SFPQ ΔN&ΔC is recruited to DNA damage sites to a significantly lesser extent.

**Figure S1.**
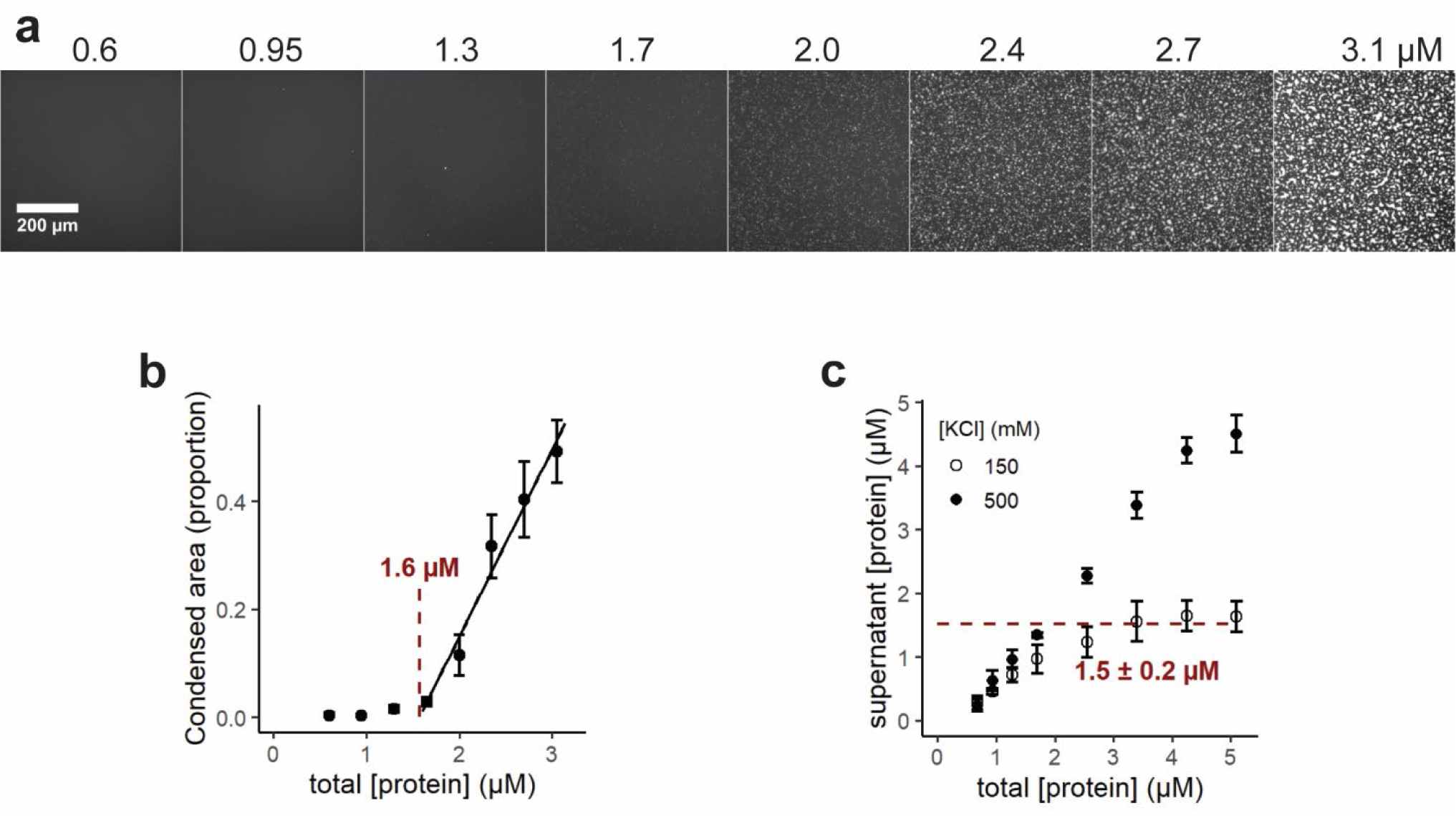
Quantification of GFP-tagged SFPQ condensed phase by fluorescent imaging confirms the *C*_sat_ determined using centrifuge-based assay. **(a)** Representative images of fluorescent mEGFP-SFPQ condensed phase on the plate surface after all droplets had settled. **(b)** Quantification of the total area covered by the condensed phase reveals a sharp increase in condensed phase after a total SFPQ concentration of 1.6 μM, consistent with **(c)** the *C*_sat_ determined by measuring the protein concentration in the dilute phase. Error bars are SD; n=3.

**Figure S2.**
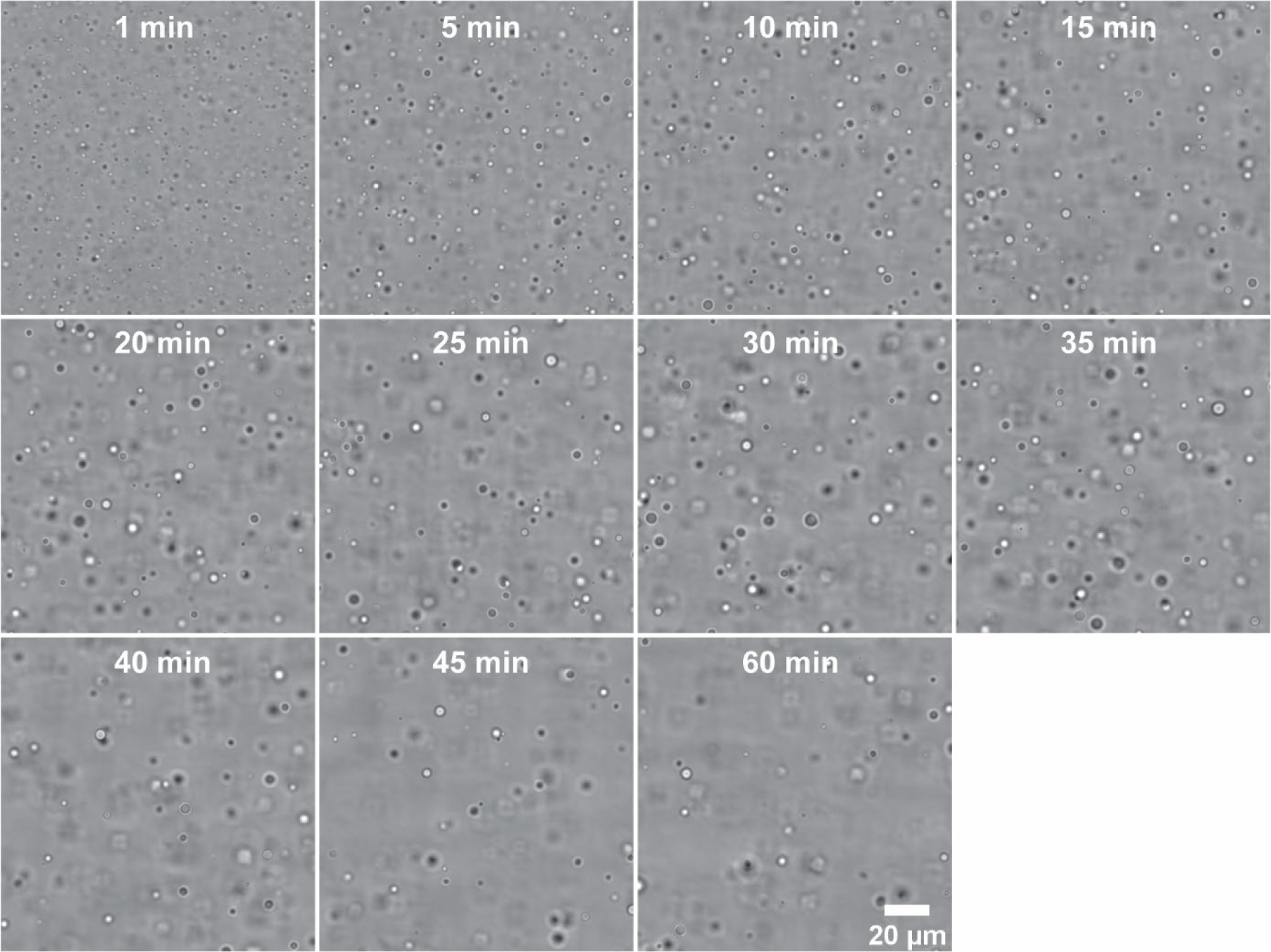
SFPQ droplets imaged at different time intervals after inducing phase separation. mEGFP-SFPQ (full-length) droplets were imaged at 5 min intervals to assess the time-dependence of droplet formation and settling. Droplet size and number appeared to stabilise after 10-15 min, before the number of droplets in suspension visibly decreased due to settling at time points after ∼ 35 min. Sample contains a total protein concentration of 5.1 μM in buffer containing 150 mM KCl.

**Figure S3.**
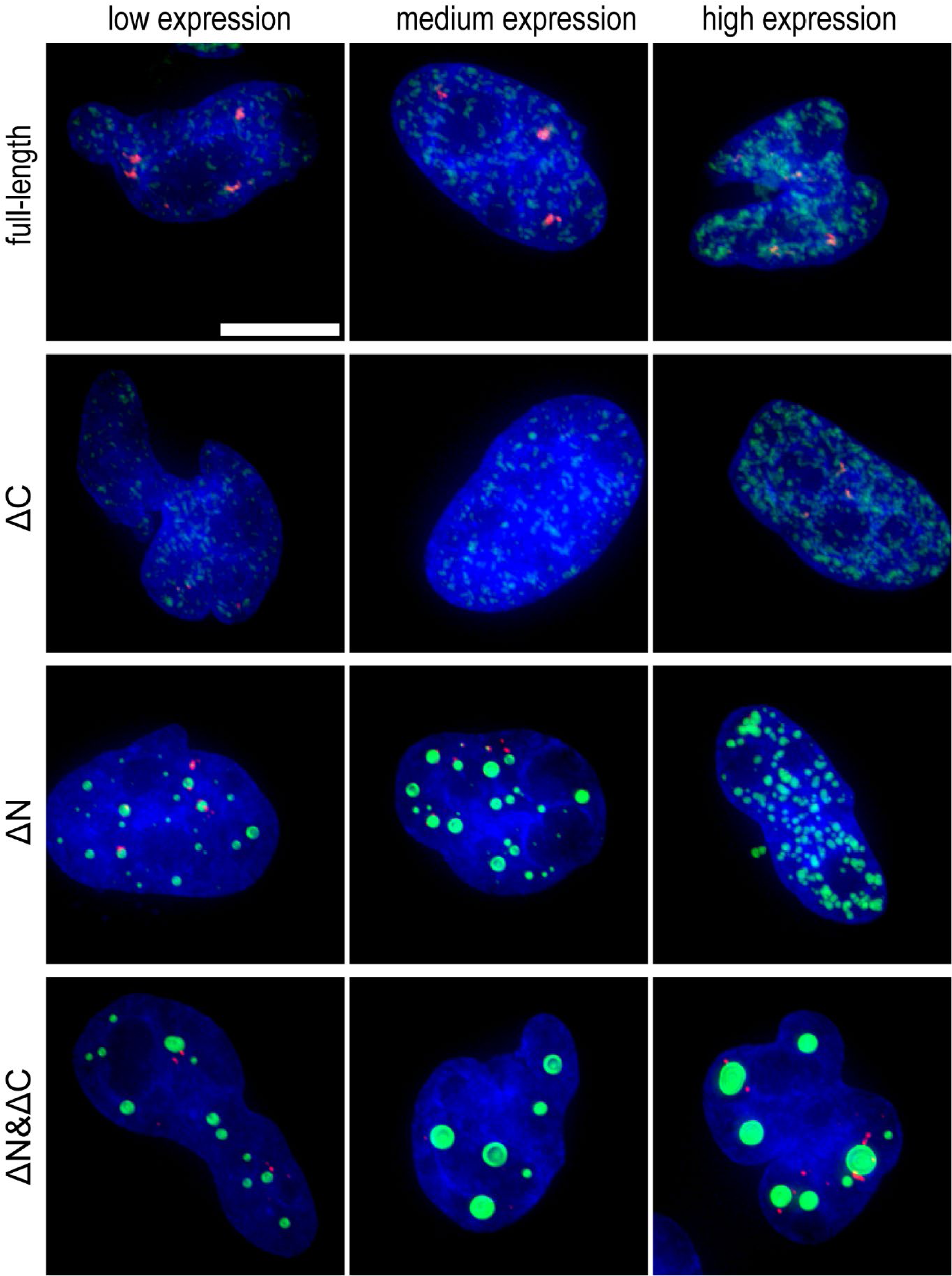
Overexpression of different GFP-tagged SFPQ truncations after transient transfection results in qualitatively similar localisation patterning in cells expressing different amounts of exogenous protein. “Low expression” (left) includes images of Hela cell nuclei for which the total integrated GFP intensity is approximately half of mean integrated intensity for all GFP-expressing nuclei in the experiment. “Medium expression” (middle) shows nuclei with integrated GFP intensity approximately equal to the mean intensity. “High expression” (right) shows nuclei with integrated GFP intensity approximately equal to double the mean intensity. Blue = DAPI; Green = GFP fluorescence; Red = NEAT1_2 FISH. Scale bar = 10 µm.

**Figure S4.**
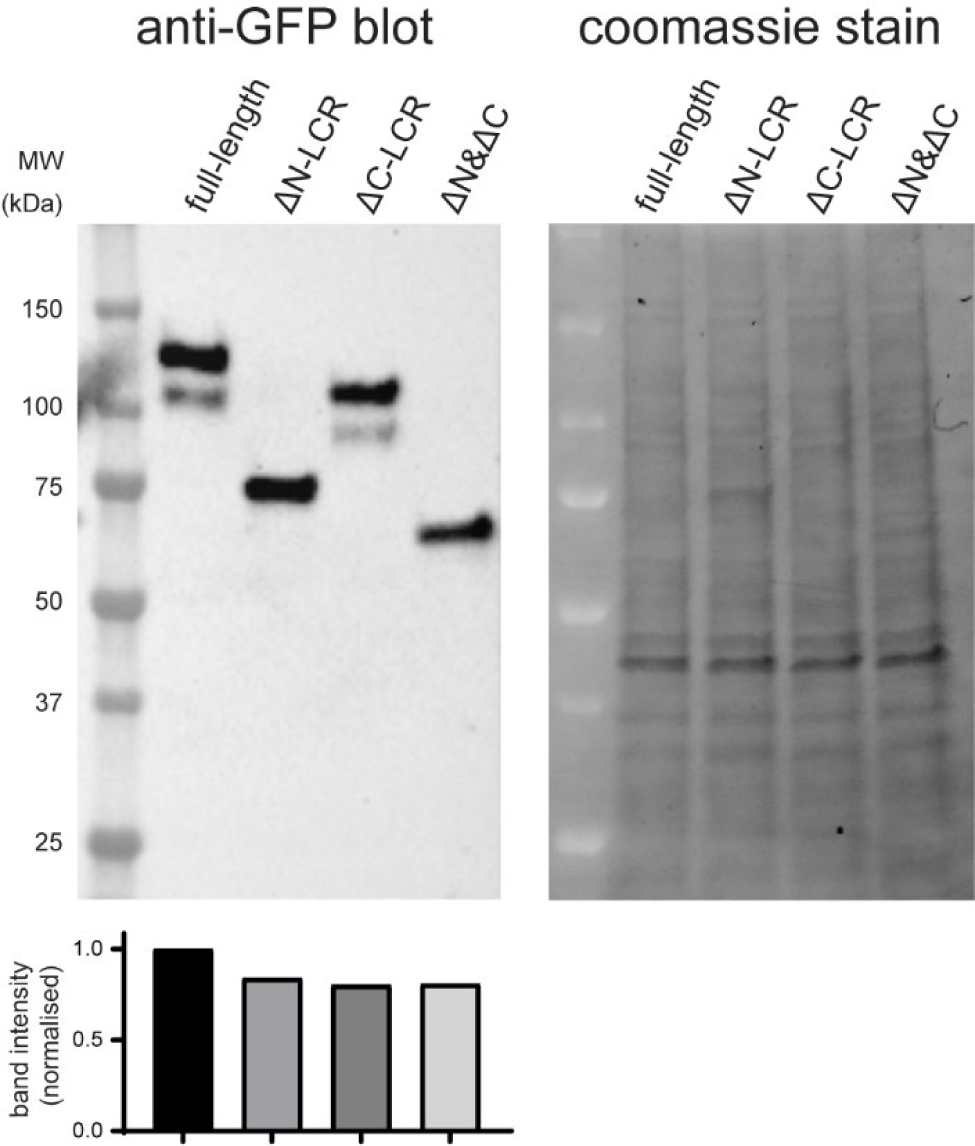
Western blot showing that average expression levels for all four GFP-SFPQ protein variants in Hela cells were similar. The gel stained with Coomassie blue (right) is shown as loading control.

**Figure S5.**
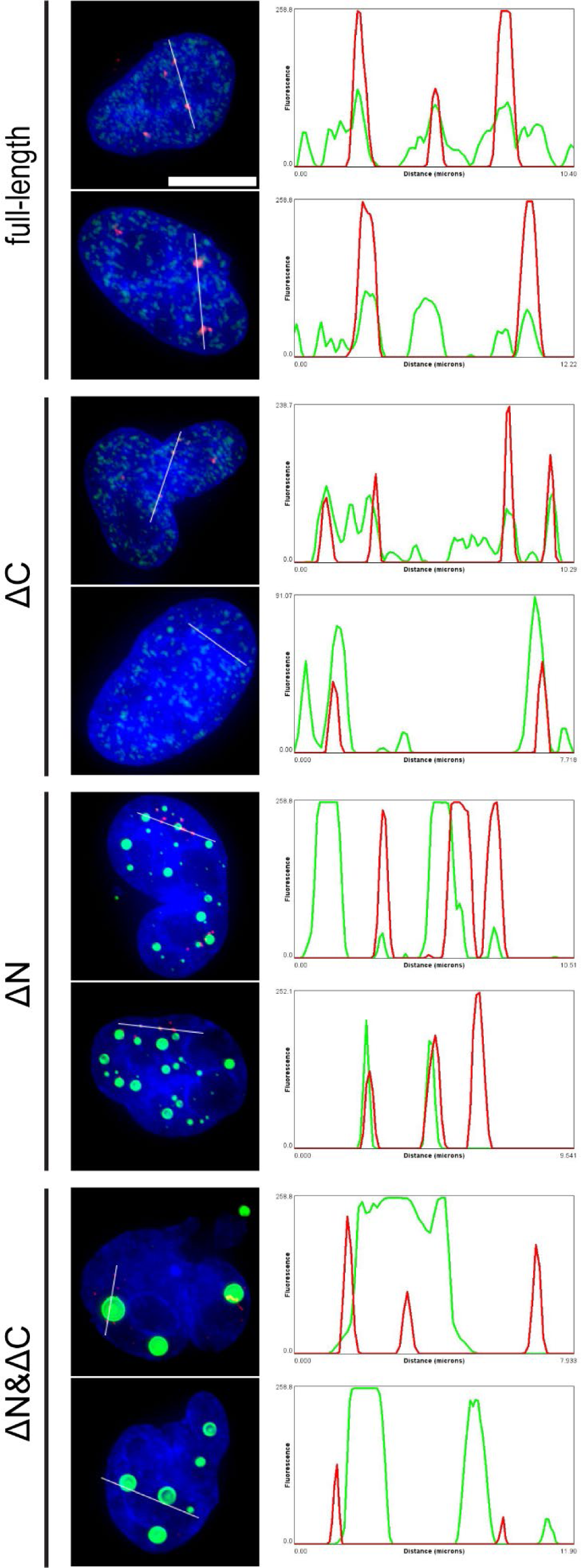
Additional linescans showing the different subnuclear localisation of GFP-SFPQ protein variants. Blue = DAPI; Green = GFP fluorescence; Red = NEAT1_2 FISH. Scale bar = 10 µm.

**Figure S6.**
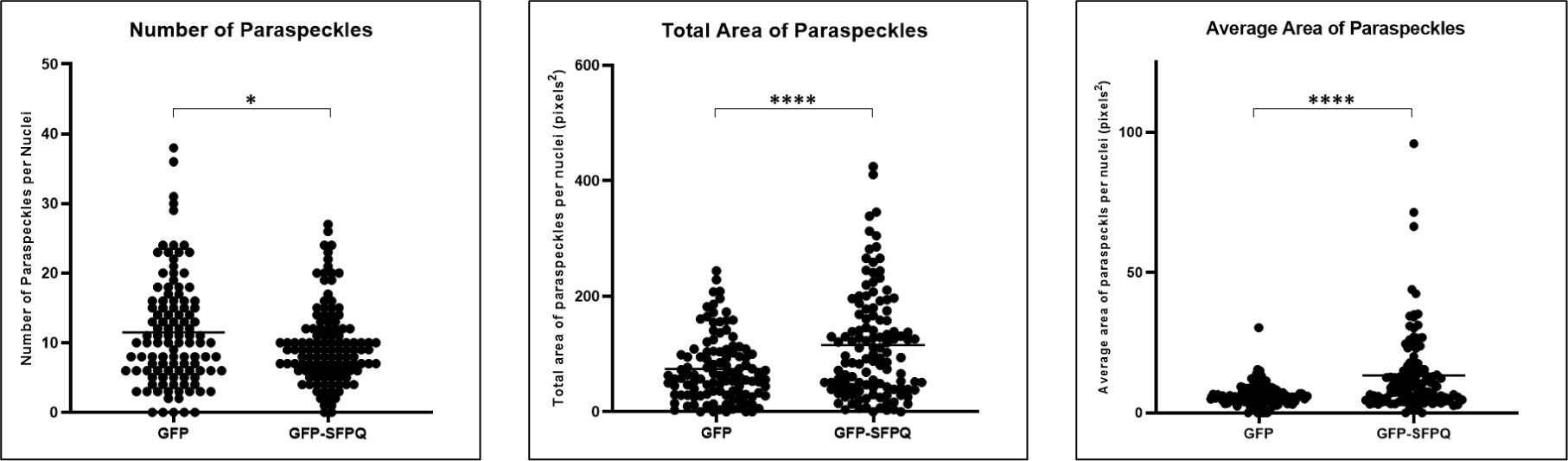
Overexpression of GFP-SFPQ increases paraspeckle area when compared to GFP overexpression (middle and right panels; **** p<0.00001 by unpaired t test with Welch’s correction). This is accompanied by a small but significant decrease in the number of paraspeckles (left panel; * p<0.01).

**Figure S7.**
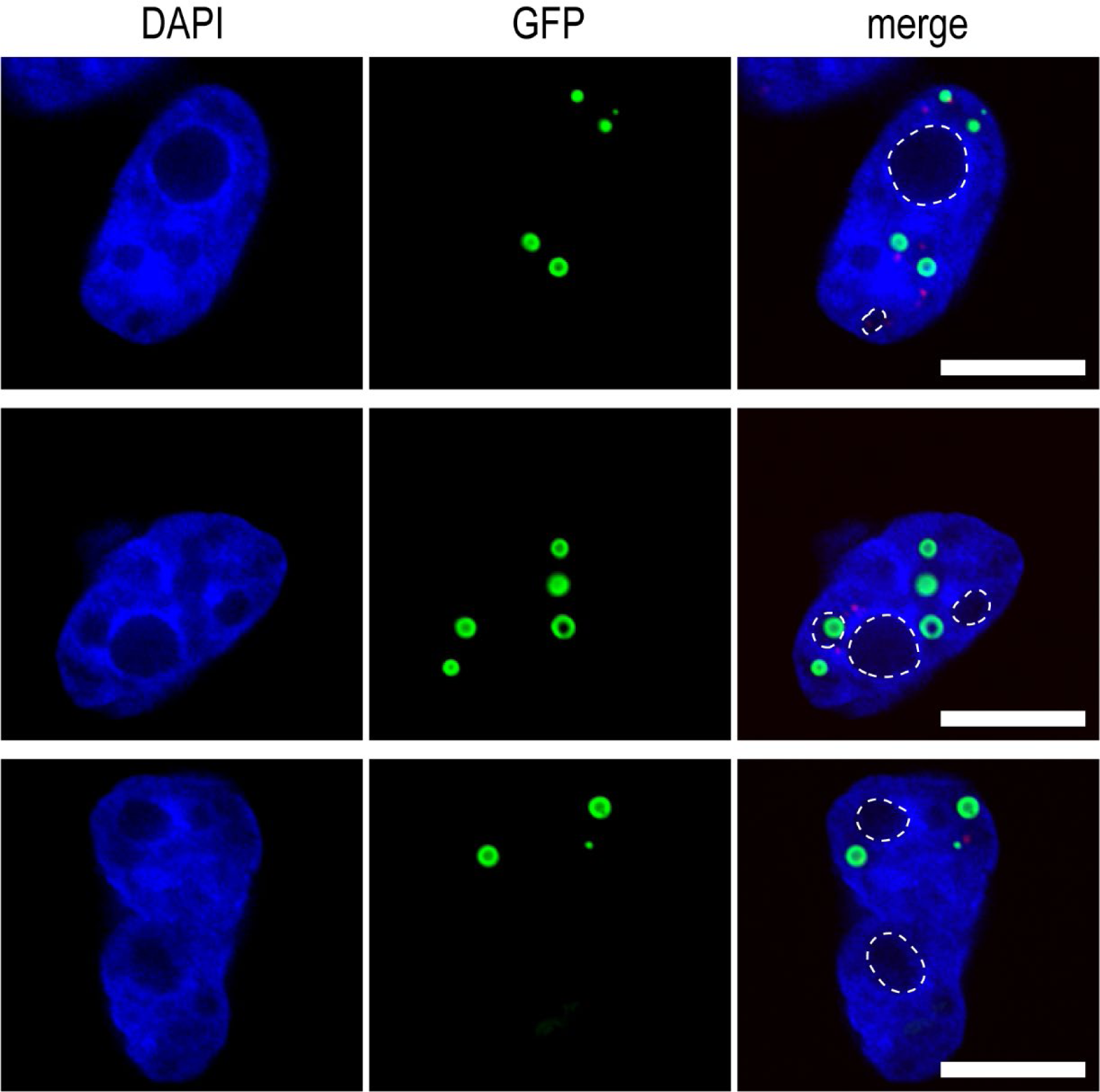
SFPQ ΔN&ΔC condensates are distinct from nucleoli. Images are single slices from the z-stacks used to construct maximum intensity projections like those shown in Figure 3a. Nucleoli (DAPI-negative regions) are outlined with white dashes. Scale bar = 10 µm.

