## Supplementary material for "Different low-complexity regions of SFPQ play distinct roles in the formation of biomolecular condensates": Movies S1-S7: SFPQ LCRs Supp Movies.pptx

### Slide 1
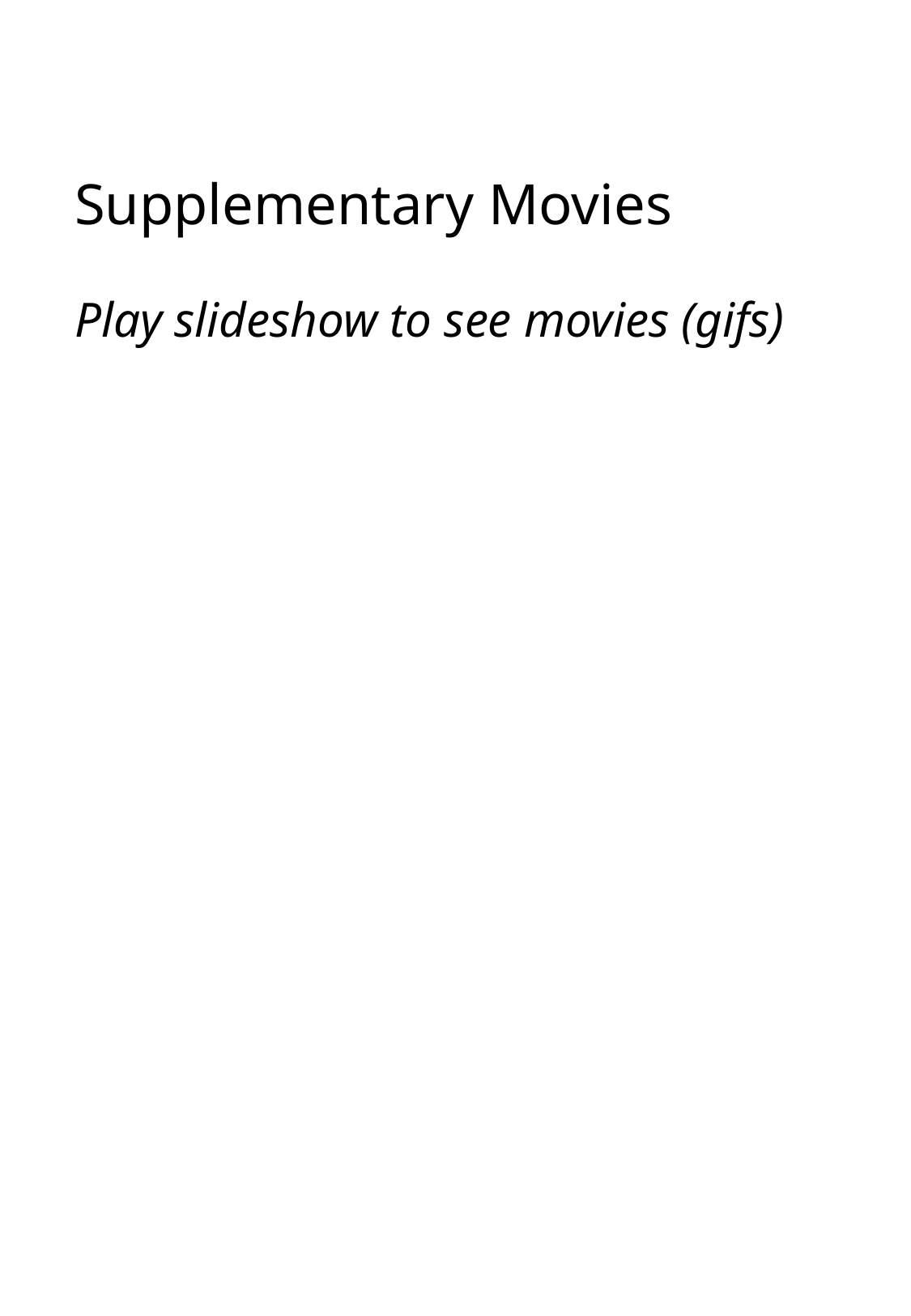

#### Supplementary Movies Play slideshow to see movies (gifs)

### Slide 2
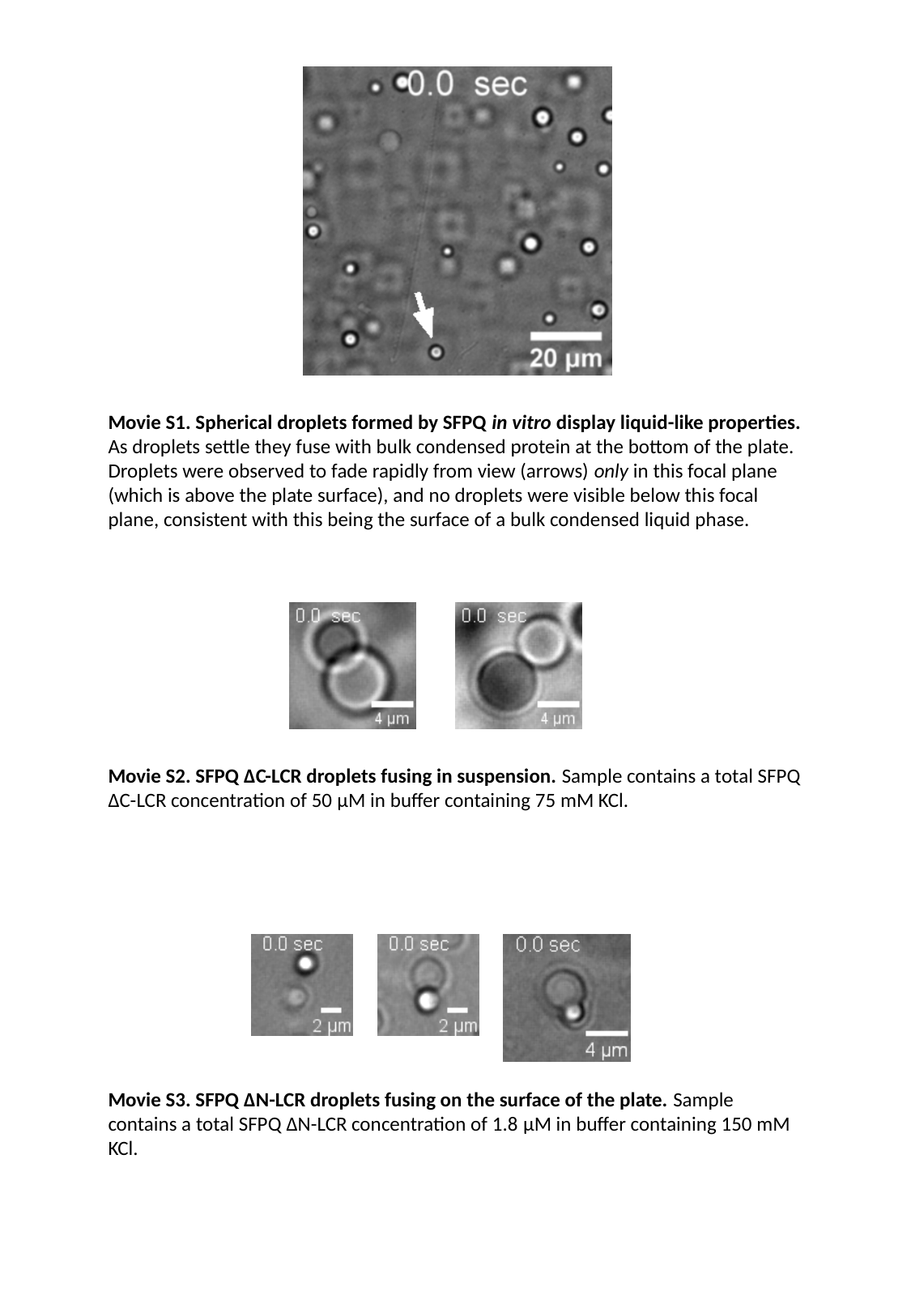

Movie S1. Spherical droplets formed by SFPQ in vitro display liquid-like properties. As droplets settle they fuse with bulk condensed protein at the bottom of the plate. Droplets were observed to fade rapidly from view (arrows) only in this focal plane (which is above the plate surface), and no droplets were visible below this focal plane, consistent with this being the surface of a bulk condensed liquid phase.
Movie S2. SFPQ ΔC-LCR droplets fusing in suspension. Sample contains a total SFPQ ΔC-LCR concentration of 50 μM in buffer containing 75 mM KCl.
Movie S3. SFPQ ΔN-LCR droplets fusing on the surface of the plate. Sample contains a total SFPQ ΔN-LCR concentration of 1.8 μM in buffer containing 150 mM KCl.

### Slide 3
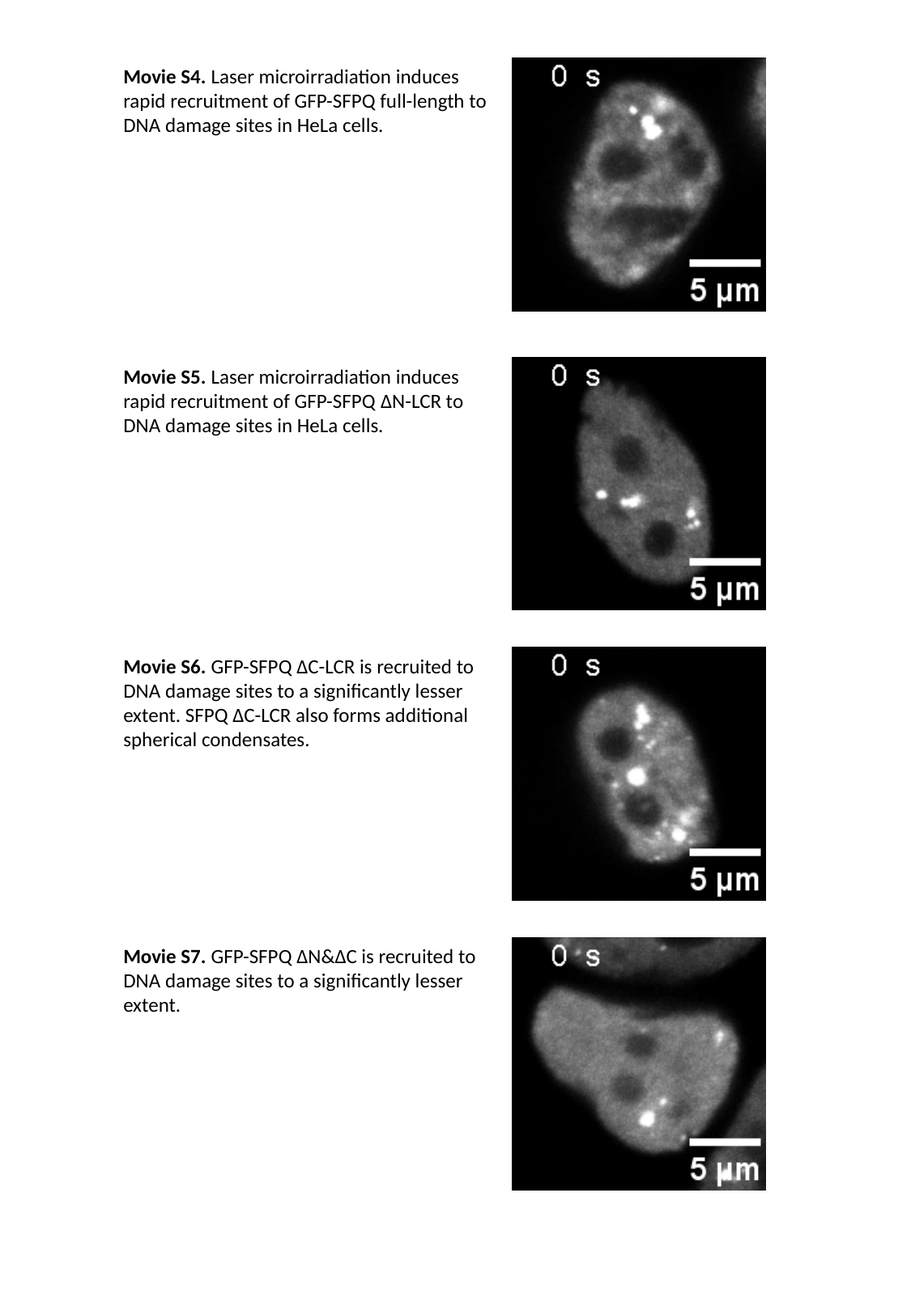

Movie S4. Laser microirradiation induces rapid recruitment of GFP-SFPQ full-length to DNA damage sites in HeLa cells.
Movie S5. Laser microirradiation induces rapid recruitment of GFP-SFPQ ΔN-LCR to DNA damage sites in HeLa cells.
Movie S6. GFP-SFPQ ΔC-LCR is recruited to DNA damage sites to a significantly lesser extent. SFPQ ΔC-LCR also forms additional spherical condensates.
Movie S7. GFP-SFPQ ΔN&ΔC is recruited to DNA damage sites to a significantly lesser extent.
